# Single-nucleus transcriptomics of wing sexual dimorphism and scale cell specialization in sulphur butterflies

**DOI:** 10.1101/2024.10.10.617718

**Authors:** Ling S. Loh, Joseph J. Hanly, Alexander Carter, Martik Chatterjee, Martina Tsimba, Donya N. Shodja, Luca Livraghi, Christopher R. Day, Robert D. Reed, W. Owen McMillan, Gregory A. Wray, Arnaud Martin

## Abstract

The evolution of sexual secondary characteristics necessitates regulatory factors that confer sexual identity to differentiating tissues and cells. In *Colias eurytheme* butterflies, males exhibit two specialized wing scale types — UV-iridescent (UVI) and lanceolate scales — absent in females and likely integral to male courtship behavior. This study investigates the regulatory mechanisms and single-nucleus transcriptomics underlying these two sexually dimorphic cell types during wing development. We show that Doublesex (Dsx) expression is itself dimorphic and required to repress the UVI cell state in females, while unexpectedly, UVI activation in males is independent from Dsx. In the melanic marginal band, Dsx is required in each sex to enforce the presence of lanceolate scales in males, and their absence in females. Single-nucleus RNAseq reveals that UV-iridescent and lanceolate scale cell precursors each show distinctive gene expression profiles at 40% of pupal development, with marker genes that include regulators of transcription, cell signaling, cytoskeletal patterning, and chitin secretion. Both male-specific cell types share a low expression of the *Bric-a-brac* (*Bab*) transcription factor, a key repressor of the UVI fate. Bab ChIP-seq profiling suggests that Bab binds the cis-regulatory regions of gene markers associated to UVI fate, including potential effector genes involved in the regulation of cytoskeletal processes and chitin secretion, and loci showing signatures of recent selective sweeps in an UVI-polymorphic population. These findings open new avenues for exploring wing patterning and scale development, shedding light on the mechanisms driving the specification of sex-specific cell states and the differentiation of specialized cell ultrastructures.

## Introduction

Animal traits involved in sexual function often differ between the males and females of a given species. The emergence of these sexual dimorphisms from a shared genome imply the existence of regulatory mechanisms that provide cells and tissues a sexual identity during development. In insects, this somatic sexual identity is primarily acquired in a cell-autonomous fashion, with each cell genotype or karyotype (sex chromosome composition) establishing somatic sex identity via one of many sex determination pathways (*1–4*). These diverse genetic cascades generally converge on a shared integrating mechanism across insects, the differential splicing of sex-specific isoforms of the *Doublesex* pre-mRNA (*Dsx*), resulting in transcription factors called DsxM in males and DsxF in females (*5–7*). Comparative studies of Dsx functions indicate that its expression is often spatially restricted to cells with dimorphic potential, including in gonadal tissues, epithelial structures, and neuronal circuits (*8–14*). DsxM and DsxF share similar occupancy profiles across the *Drosophila* genome, suggesting their binding targets are not sex-specific (*15*, *16*). Instead, DsxM and DsxF mediate distinct, and sometimes opposite *cis-*regulatory effects to bias gene expression. For example, in the *Drosophila melanogaster* abdomen, DsxM represses the BTB-domain transcription factor gene *Bric-a-brac* (*Bab*), while DsxF activates it (*17*, *18*). As Bab represses dark melanization in the fly abdomen, this dual regulation underlies the dichromatism of *D. melanogaster* where only males are fully pigmented in their last two abdominal segments.

To understand how somatic cell identity is integrated into dimorphic gene expression programs, we can leverage the study of scale cell precursors that underlie sexual dichromatism in butterflies. Each color scale is the extension of a single cell, and each cell integrates spatial and sexual cues during development to differentiate into the biological pixels that form wing patterns. The Orange Sulphur butterfly *Colias eurytheme* is an emerging model system for the study of scale dimorphisms (*19–23*). Not only do the males and females of this species show distinct melanic patterns on their dorsal wing, but males also exhibit a bright ultraviolet-iridescent (UVI) pattern that is used as a species recognition signal by conspecific females to avoid interspecific matings (*24–27*). UV-iridescence is conferred by the specialization of their dorsal cover scales, which form dense stacks of 7-9 air-chitin layers on their upper surface in males. Females lack UVI scales and instead show typical orange scales, which do not elaborate a multilayered ultrastructure. The butterfly ortholog of *Bab* is expressed in most orange-yellow scales, where it is necessary to repress the UVI fate in all wing scale cell precursors (*28*). *Bab* is specifically silenced in the male dorsal cover scales starting at 35% of pupal development, thus activating the UVI fate by de-repression. Hybrid males carrying at least one *Bab* allele from *Colias philodice,* a monomorphic, non-UVI species, express Bab in their dorsal cover scales and lack UVI scales. In summary, Bab is both necessary and sufficient to block the terminal differentiation of UVI scales, and *C. eurytheme* alleles of Bab turn off its expression in male dorsal cover scales, thus activating the UVI state by derepression.

To understand the mechanism of sexual differentiation in the butterfly wing, it is also important to place scale differentiation into a broader developmental context. The pupal wing of lepidopteran insects consists of an epithelial bilayer, where dorsal and ventral surfaces are separated by an acellular baso-lateral membrane. The structural integrity of this tissue is maintained by small columnar epithelial cells. Scales are macrochaetes that derive from the differentiation of a Scale Organ Precursor cell (SOP) lineage, akin to mechanosensory bristles (*29*, *30*), that abrogate the forming of associated neurons and glia to only give rise to a trichogen (scale) cell precursor and an associated tormogen (socket) cell (*31–34*). Scale cell precursors become large, polyploid, and aligned along tightly arranged rows that alternate between cover scales, that will give rise to scales on the top surface, and a lower layer of ground scales (*35–37*). Pupal wings are also infiltrated by trachea, including major branches and numerous small tracheoles that transiently invade the wing epithelium during early pupal development (*38*, *39*). The major branches develop inside the lumen of epithelial tubes that persist into sclerotized wing veins, which provide robustness to the adult wing and act as a hemolymph circulatory system (*40*). Finally the pupal wing is embedded with mobile hemocytes (*38*, *39*), and likely contains a small population of neuroglial cells due to the presence of mechanosensory and thermosensitive sensillae, particularly along the wing veins (*40*).

Here we delve into the cell type diversity of *C. eurytheme* developing wings with the overarching goal to link gene expression programs and the specific differentiation of complex scale ultrastructures, such as the UV-iridescent scales found in males. First, we test the effect of *Dsx* loss-of-function in wing development in both sexes to infer its roles in sexual dimorphism. Second, we analyze single-nucleus transcriptomes of the male pupal wing and profile gene expression in scale cell precursors at a cellular resolution, with a focus on differentiated clusters that correspond to two male-specific, specialized cell types. Last, we integrate transcriptome signatures with ChIP-seq profiles of Bab occupancy to sketch a set of potential Bab transcriptional targets involved in UV-iridescent scale differentiation. Overall, this study illustrates how color scale types emerge from divergent gene expression programs, and provides foundational knowledge for the study of cell sexual dimorphism and ultrastructural specialization.

## Results

### Two male-specific scale types with divergent ultrastructures

The sexual dimorphism of *C. eurytheme* butterflies manifests on the dorsal surface of their wings, where males show bright UV iridescence across its medial orange region, as well as a thin, continuous marginal black band (**Fig. 1A**). There is no UV iridescence in wild-type females, and their marginal patterns are wider, jagged at their interface with the orange area, and flecked by yellow spots (**Fig. 1A**). The two sexes thus differ in UV iridescence as well as in the shape of the marginal band. We used Scanning Electron Microscopy to further survey the ultrastructure of scale types in *C. eurytheme* (**Fig. 1B-H**). Typical scales that are found on either sex or wing surface share a 2 μm distance between their apical ridges (2.0± 0.1 μm, N = 125 measured scales), a feature that is relatively constant across melanic scales (*e.g.* forewing discal spots of both sexes), pterin-pigmented scales (orange, yellow), and pterin-deficient scales (*e.g.* white scales from Alba female morphs). In contrast, UVI scales are not only characterized by the multilayering of their ridge lamellae (*20*, *26*, *41*), but also by the density of the ridges themselves, with a distance between ridges averaging only 1 μm (1.0± 0.1 μm, N = 25 scales) (**Fig 1D, G**). A second type of male-specific scales type, here dubbed the lanceolate scales, show a large inter-ridge distance of 4.4 ± 0.5 μm (N = 20 scales) that gives them a corrugated look under light microscopy. These scales adorn an unusual ultrastructure, with crossrib structures that resemble soybean pods joining the longitudinal ridges (**Fig. 1D, H**), and that overlay a porous inner matrix (*42*). It was proposed that these scales play a role in pheromone retention or spreading, due to their male-specificity and the spongy aspect of their ultrastructure (*24*, *43*, *44*). Lanceolate scales occur specifically on the cover scale layer of the male melanic bands, where their wide apical lobes overlay a layer of yellow ground cover scales. In females, the dorsal marginal band is devoid of lanceolate scales and shows instead canonical melanic scales as cover scales. In summary (**Fig. 1I-J**), the dorsal surface of *C. eurytheme* wings includes two male-specific scale types that each feature unique ultrastructures – the UVI scale and lanceolate scales.

**Figure 1.**
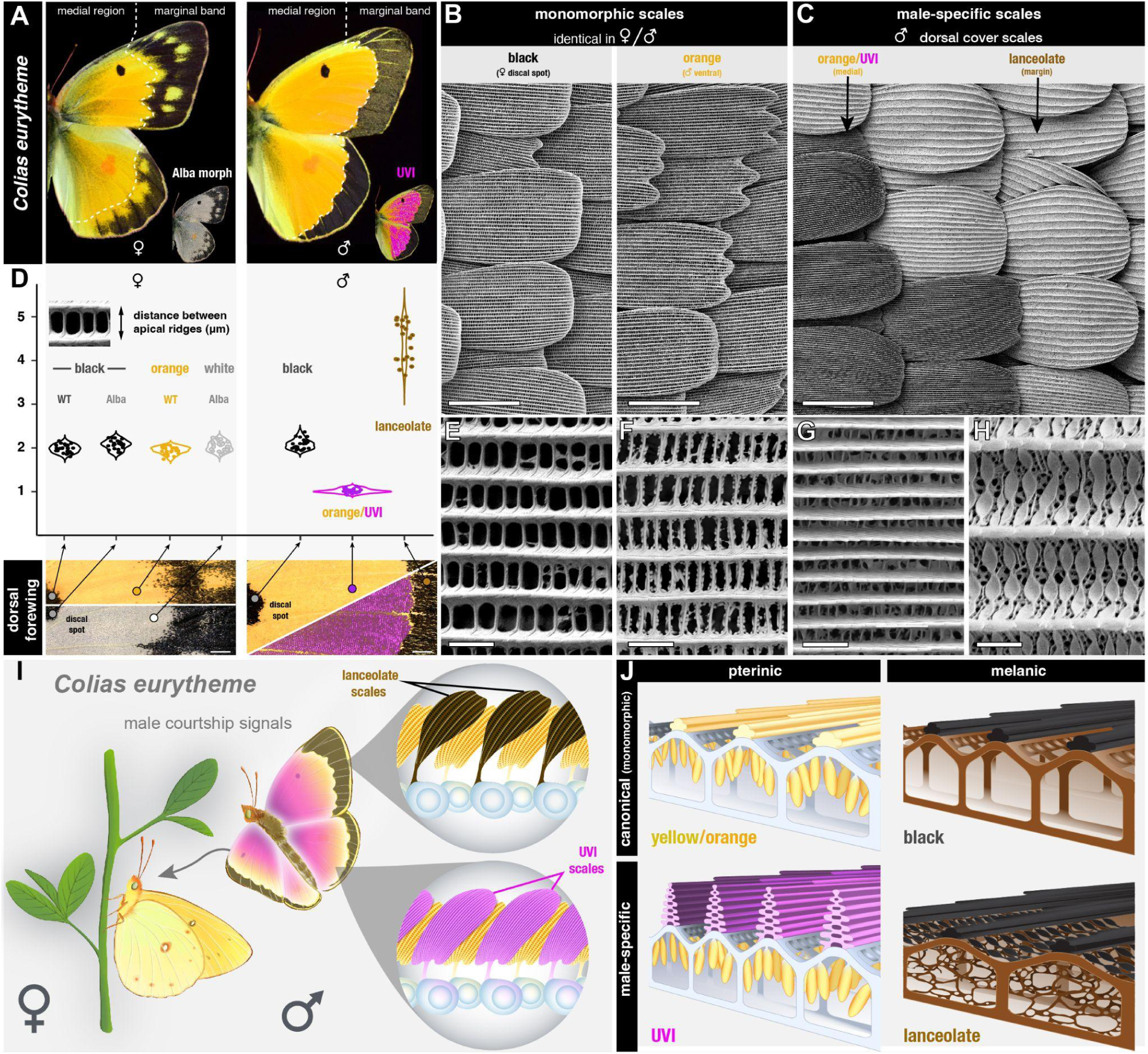
Male-specific scales in the dorsal wings of *C. eurytheme*. **A.** Dorsal views of male and female butterflies, highlighting the difference in width and patterning of the marginal bands, and the male-specific UV-iridescence (magenta ; an UV-photograph was colored in magenta and overlaid on a normal image). Alba morphs are a female-limited phenotype characterized by pterin-pigment deficiency in their scales. **B-C.** Scanning electron micrographs of representative non-dimorphic scales (here in B, left : female dorsal forewing melanic scales of the discal spot ; right : male ventral orange scales), and male-specific scales (here in C: interface between UVI and lanceolate scales, dorsal hindwing). **D.** Mean apical ridge distance in individual scales from melanic discal spots, medial regions, or male marginal bands, highlighting the derived ultrastructure of males-specific UVI and lanceolate scales. **E-H.** Apical views of the ultrastructure of representative scale types (same scale types as in panels B-C), featuring the longitudinal ridges (horizontal structures) and the transversal microribs (vertical). **E**: female dorsal cover black scale; **F**: male ventral cover scale (orange, non-UVI); **G**: male UVI dorsal cover orange scale; **H**: male lanceolate dorsal cover margin scale. **I.** Position of male-specific cell types on the dorsal wings. The cell bodies of scale precursors (grey) are transient structures that do not occur in adults. **J.** Schematic representation of the ultrastructures (viewed as transversal cross-sections) from the four main scale types – canonical (orange and black), UVI, and lanceolate. Scale bars; B-C = 50 μm; D = 1 mm ; E-H = 2 μm.

### *Doublesex* has distinct functions on marginal patterning and UV dichromatism

Gene loss-of-function assays targeting *Doublesex* generate intersexual phenotypes across insects, with secondary sexual characteristics losing their sex-specific differentiation, and sometimes reversing to the state found in the opposite sex (*9*, *45–50*). We thus used CRISPR targeted mutagenesis to generate G_0_ adults carrying *Dsx* mosaic knock-outs (abbreviated mKO, or “crispants”), using single sgRNAs that targeted either the DNA-binding Domain or the Dimerization Domain shared across *Dsx* isoforms (*6*), resulting in deletion alleles at the target sites (**Fig. S1)**. A total of 33 crispant adults showed mosaic effects on sexually dimorphic features of the dorsal wings (**Fig. 2**, **Fig. S2-3, Table S1**). *Dsx* crispants from both sexes showed intermediate intersexual states in the aspect of the marginal bands, the melanic patterns at the distal edge of their wings. The marginal bands of female crispants showed a narrowing of the melanic marginal band compared to wild-type females, resulting in a loss of yellow spots. The reverse was observed in the marginal bands from male crispants, with a partial expansion of the melanic band resembling the female state.

**Figure 2.**
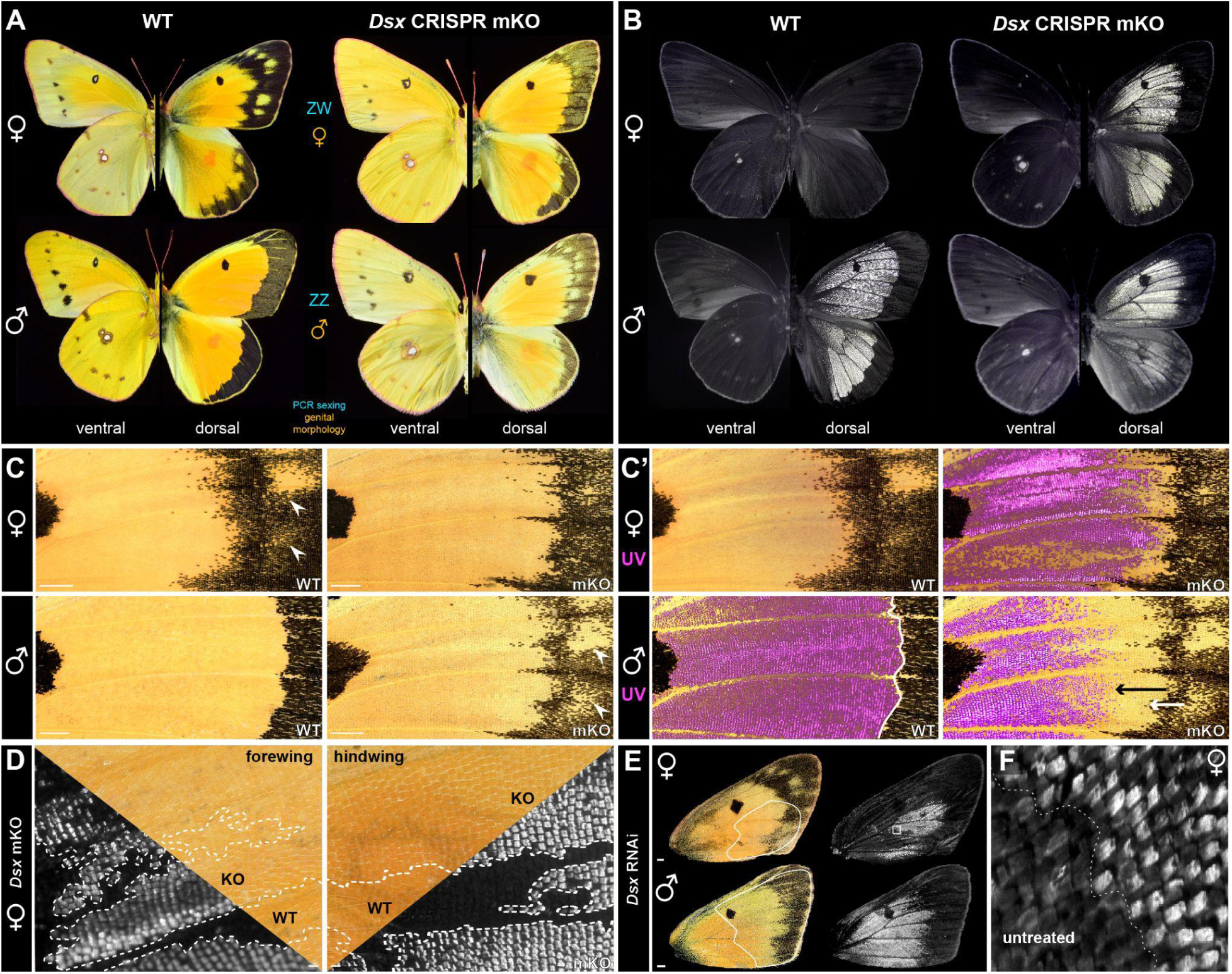
*DsxF* blocks UV-iridescence in females in a cell-autonomous fashion. **A.** Effects of *Dsx* mosaic knock-outs (mKO), as seen in comparisons of female and male WT individuals with representative *Dsx* G_0_ crispants. The sex of each crispant was determined by concordant genital morphology and genotyping. Phenotypic effects are limited to the dorsal surfaces and most visible in the marginal region. **B.** UV-photography (320-400 nm) of UV-iridescent dorsal patterns in the same mutants, showing an ectopic gain of UVI scales in females (top). In males (bottom), the feminization of the marginal region triggers a regression of the male UVI pattern distal border. **C-C’** Close-up views of the central forewing regions (same individuals as in panels A-B), including overlays of UV photographs (C’: magenta false-color) over visible light images, and highlighting the intermediate states of marginal patterns. In females : gain of UVI scales, regression of the melanic marginal patterns. In males : gain of female yellow spots in males (arrowheads), extension of the melanic marginal patterns (white arrow), and regression of the UVI field (black arrow). **D.** Cell autonomy of UVI scale gains in female *Dsx* mKOs, as shown by continuous UVI mutant clones with sharp boundaries (KO). Superimposed views of the same region, taken in visible and ultraviolet light, are shown across each diagonal line. **E.** RNA interference effects of *Dsx* siRNA electroporation in male and female forewings. Dotted lines mark the approximate areas that were electroporated. *Dsx* knockdown results in ectopic UVI (top), and in a feminization of the marginal pattern in males, including with a regression of medial UV iridescence at its distal border. **F.** High-magnification view of the female wing shown in panel E, at the interface of the treated (ectopic UVI scales) and untreated area. Scale bars: C, E = 1 mm; D = 100 μm.

The effects of *Dsx* mKOs on UV iridescence (UVI) indicated two distinct categories of effects between males and females. Female crispants showed a gain of UVI that phenocopies the male state (**Fig. 2A-C’**). This conversion towards UVI scale types occurred along sharp clone boundaries (**Fig. 2D**), showing that DsxF is necessary for the repression of UVI in cell autonomous fashion. In contrast, while male crispants show a spatial reduction of the UVI field (**Fig. 2A-B**), this corresponds to a non-cell autonomous shift in the positioning of its distal boundary, in the vicinity of the marginal region (**Fig. 2C-C’**). Indeed, none of the male crispants showed a loss of UVI in the proximal and central sections of the wing that would indicate cell-autonomous effects (**Fig. S2**). In addition, we electroporated short-interfering RNAs (siRNAs) to drive RNAi knockdowns of *Dsx* at the pupal stage (*10*, *51*). UVI scales were unaffected throughout the central domain of the RNAi treated male dorsal forewings (**Fig. 2E**). Consistently with the mosaic knock-outs, female wings electroporated with *Dsx* siRNA showed ectopic UV iridescence in the cover scales (**Fig. 2E-F**). To explain these non-cell autonomous effects in males, we extrapolate that the spatial patterning of the marginal region includes sex-specific inputs on morphogenetic signaling events, that in turn determine its width and sub-division into fields of melanic *vs.* non-UVI yellow scales.

### DsxF and Bab activation repress UV iridescence in female dorsal cover scales

Both *Dsx* KOs and knock-downs result in ectopic UV iridescence across the female medial region, and conversions to the UVI scale type are restricted to cover scales (**Fig. 3A-B**). This cover scale specificity is unlike the effect of *Bab* KOs (*28*), which converts the both ground and cover scale layers to an UVI state. Next, we profiled the expression of Dsx between 30-40% of pupal development – a temporal window where Bab becomes downregulated in male UVI scale cells. To do this, we use a monoclonal antibody that recognizes both the Dsx-DBD (DNA-binding domain) shared by both isoforms (*52–54*). Because Dsx isoforms are sex-specific in Lepidoptera, for simplicity we call the detected antigen DsxF or DsxM based on the sex of the dissected pupae. In females, DsxF is detected throughout both the medial and marginal region with a strong enrichment in the dorsal cover scales (**Fig. 3C**), while Bab is expressed in both female ground and cover scales (**Fig. 3C’**). In contrast, DsxM is restricted to the dorsal cover scales of the wing margin in male wings (**Fig. 3D-D’**). It follows that the lack of cell-autonomous effect of *DsxM* perturbation on UVI scales is likely due to its lack of expression in the medial region at this stage.

**Figure 3.**
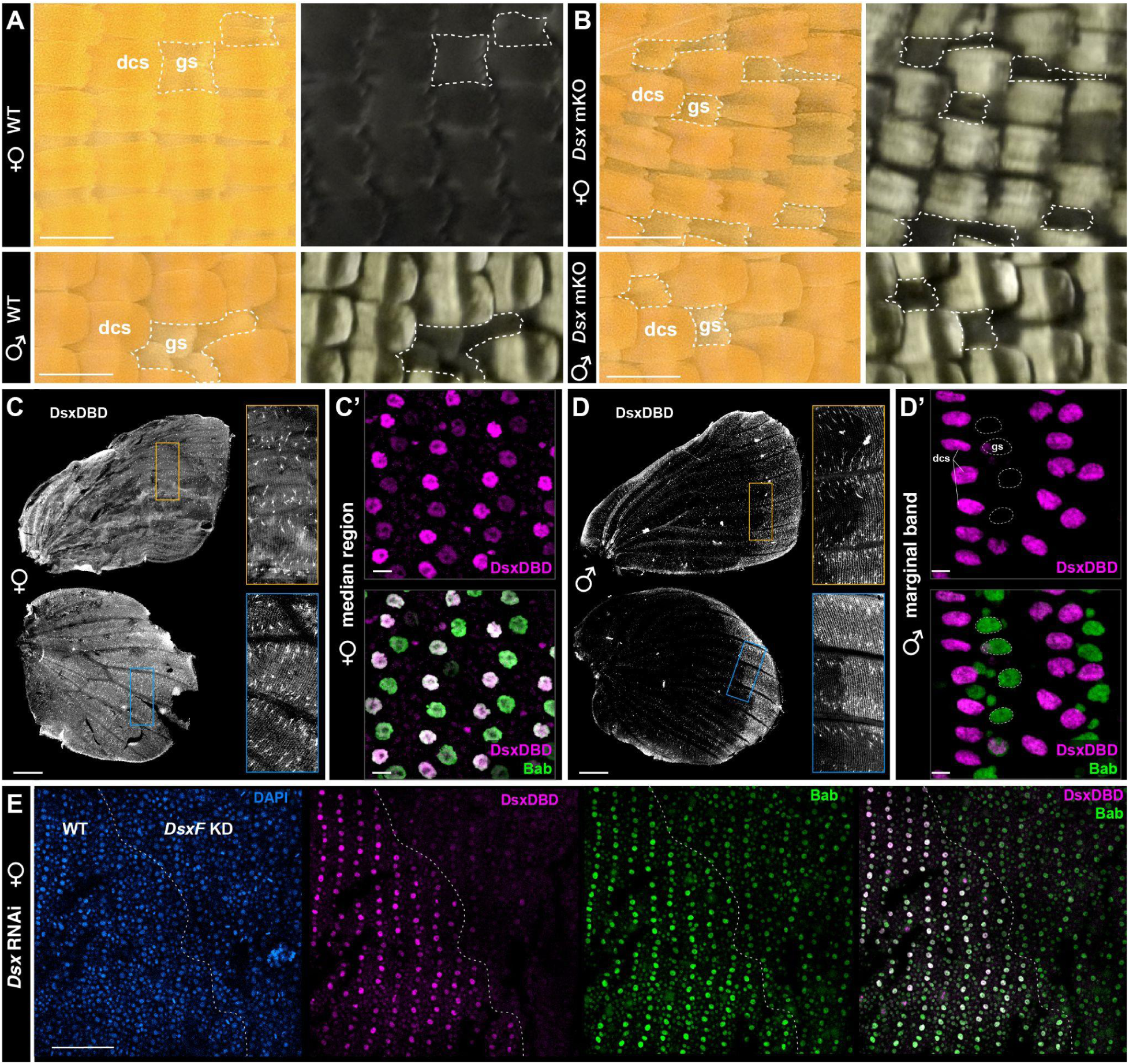
Sexually dimorphic expression of Dsx activates Bab in the female dorsal cover scales. **A-B.** High-magnification views of central dorsal wings, showing dorsal cover-scale (dcs) specific transformation of non-UVI to UVI scales in female *Dsx* KOs. Males are unaffected by *Dsx* mKOs in the central region. Dashed lines indicate small patches of the wing where ground scales (gs) are exposed. **C-D’**. Immunofluorescent detection of the DsxDBD antigen in the dorsal layer of female and male wings between 30-40% of development (shown here, C, D-D’ : 40% stage ; C’ : 30%). The Bab antigen (green) marks both dorsal cover scales (dsc) and ground scales (gs) in females (C’), and only the ground scales in males. **E.** Reduced Bab detection in the medial region of a female forewing electroporated with *Dsx* siRNAs, shown at the 30% pupal stage. Electroporation was directed to a small area controlled by the position of a conductive buffer droplet, and the dashed line marks the boundary between high and low Dsx antigen detection, corresponding to the electroporated (*DsxF* knockdown, KD) and untreated (WT) areas of the wing. Scale bars: A-B, E = 100 μm; C, D = 1 mm ; C’, D’ = 10 μm.

### Dsx controls the sex-dependent identity of margin cover scales

While the expression of Dsx is dimorphic in the medial region, it is similar between males and females in the marginal band, where it marks dorsal cover scales. Marginal dorsal scale cells are sexually dimorphic, with a canonical melanic type in females vs. the derived lanceolate type in males. Accordingly, *Dsx* CRISPR KOs showed complete reciprocal transformation of the melanic dorsal cover scales between these marginal/dorsal scale types between sexes, from typical melanic to lanceolate in *DsxF* mKOs, and vice-versa from lanceolate to typical in *DsxM* crispants (**Fig. 4A-C**). Thus, while only DsxF is required for specifying the UV dichromatism, both Dsx isoforms are required to instruct the correct patterning of marginal patterns and the identity of male-specific scales within them.

**Figure 4.**
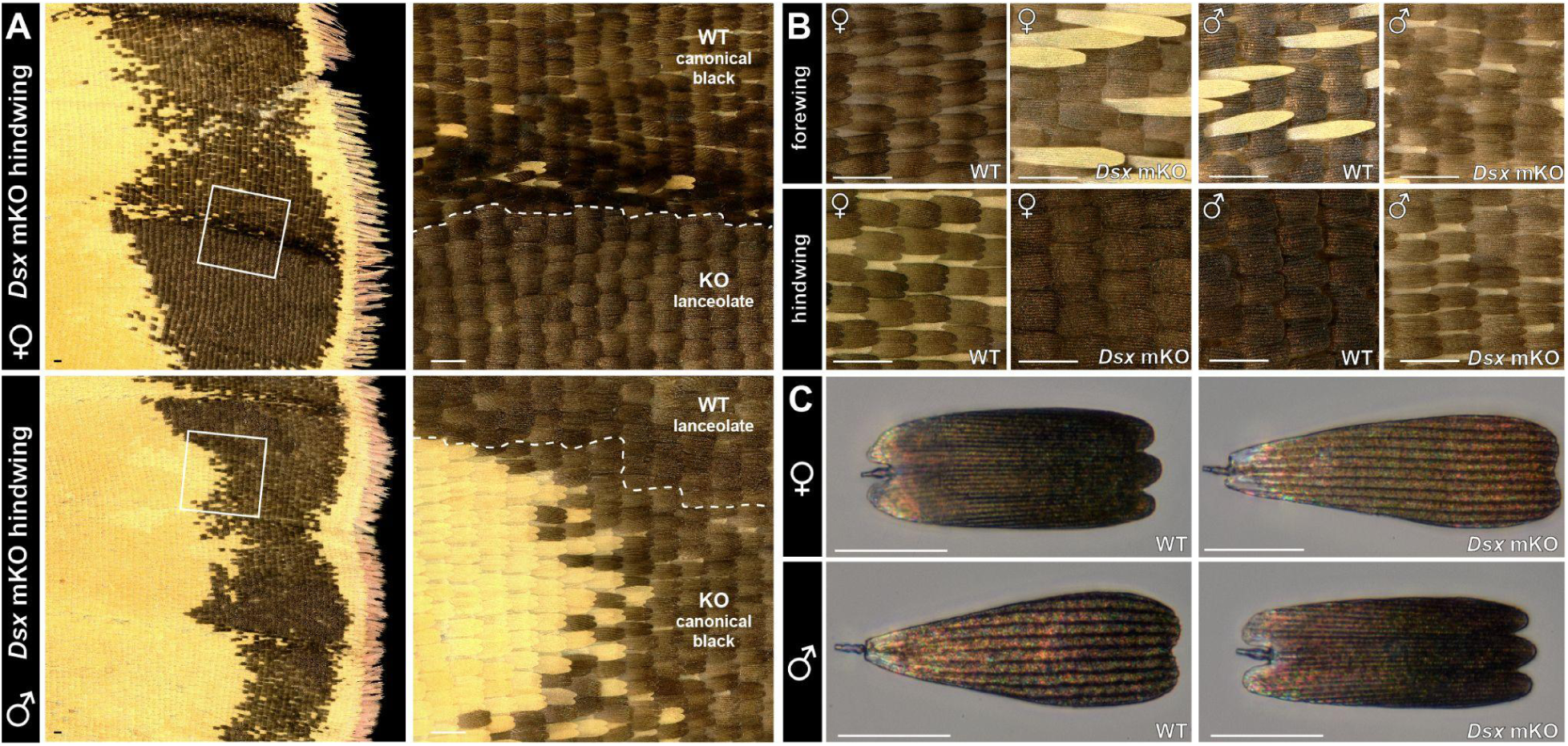
Doublesex controls the specification of male-specific lanceolate scales. **A.** Magnified views of the dorsal hindwing marginal bands of mosaic *Dsx* G_0_ crispants. KO clone boundaries are visible with yellow/orange color shifts in the medial region, and in the marginal band, with shifts from canonical black scales to the lanceolate scales in females, and vice-versa in males. **B-C.** Complete, reciprocal shifts in scale composition (B) and melanic scale identities (C) in both female and male Dsx crispants. Scale bars: A-C = 100 μm.

### The combinatorial logic of dimorphic scale type specification

We summarize below how Dsx controls multiple aspects of sexual dimorphism on the dorsal wing (**Fig. 5A**). The spatial expression of Dsx is sexually dimorphic at the 30-40% pupal stage : female wings express it in all the dorsal cover scales, while males only express it in the dorsal cover scales of the marginal region. Dsx influences the patterning of the marginal band via non-cell autonomous effects, determining the spatial extent of the melanic band and of non-UVI yellow outlines and spots. The differences in spatial Dsx expression likely explain the asymmetric effects of its perturbation in each sex. In the medial region. DsxF is required to repress the differentiation of dorsal cover scales into the UVI scale type. In the wing margin, sex-specific Dsx isoforms are required to specify where dorsal cover scales differentiate into male lanceolate scales, or into canonical melanic scales. Dsx and Bab are thus two key determinants of sexually dimorphic scale determination. We therefore sought to develop a phenomenological model that integrates the effects of both genetic switches on male-specific cell type specification.

**Figure 5.**
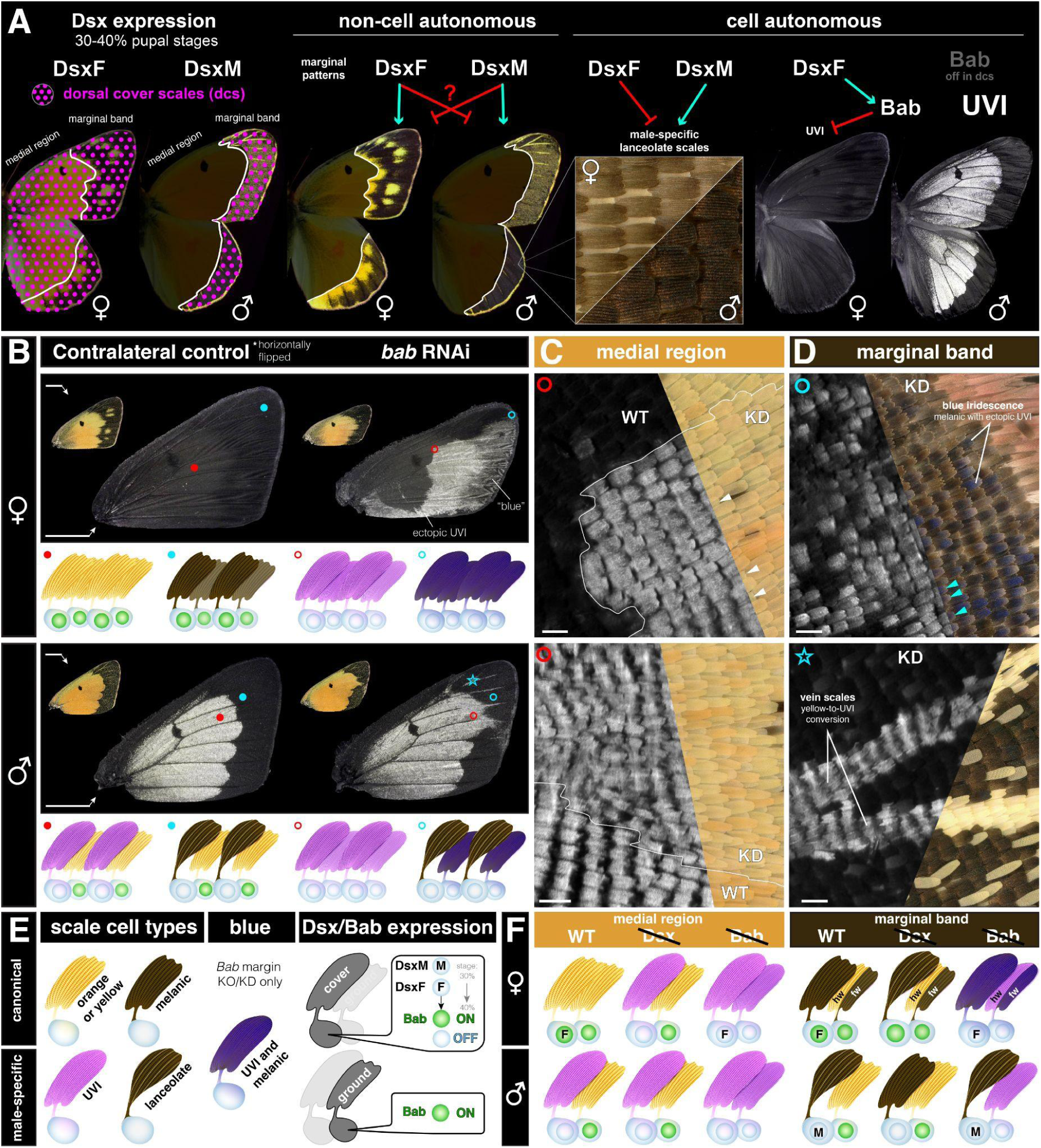
Dsx and Bab control sexually dimorphic scale fates. **A.** Working hypothesis for the effects of *Dsx* on sexually dimorphic wing traits, including non-cell autonomous patterning effects on marginal patterns, cell-autonomous requirements in the specification of lanceolate *vs.* canonical margin scales, and female-specific repression of UV-iridescence in dorsal cover scales. **B-D.** *Bab* RNAi knockdowns on the dorsal forewing phenocopy mosaic KO effects (*28*). Bab-expressing cells acquire an UVI state upon *Bab* perturbation, with the exception of the lanceolate scales, that are unaffected. This includes canonical melanic scales that acquire an UVI ultrastructure while maintaining melanism, resulting in a dark blue iridescent phenotype (D, cyan arrowheads). Ground scales are transformed to yellow UVI scales in each sex (*e.g.* in the medial region : C, white arrowheads), except in the female forewing marginal band where they convert from melanic to blue iridescent (combination of melanic and UVI features). Star : inset features a yellow-scaled veins (UV-negative in WT and controls) that acquire an UVI fate upon Bab RNAi, indicating successful knockdown in this area. **E-F.** Summary of Dsx and Bab expression and perturbation assays in dorsal wing surfaces. Scale bars: B = 5 mm ; C-D = 100 μm.

We replicated the effects of *Bab* mKOs (*26*) using RNAi knockdowns targeting the dorsal surface of the forewings from each sex, and summarized the observations from both sets of perturbation experiments (**Fig. 5B-D**). In females, Bab-deficient cells all acquire an UVI-scale morphology including UV-iridescence and high ridge density. Because all the female scale cell precursors express Bab, this is visible in both cover and ground scales of both the medial and marginal regions. In the marginal region, UVI scales retain a melanic state, acquiring a dark blue iridescence that is visible with the naked eye when illuminated with a low angle of incidence. In males, only ground scales express Bab, and according, *Bab* perturbation results in ground scale specific conversions to UVI states.

Together, these data suggest a simple combinatorial logic for male-specific cell type specification (**Fig. 5E-F**). Across the entire dorsal wing, ground scales express Bab and repress UVI without Dsx input. In the marginal cover scales, the expression of DsxF vs DsxM determines non-lanceolate vs lanceolate states. In the medial cover scales, Dsx is expressed in a female-specific fashion (as DsxF) and required for the Bab-dependent repression of UVI states.

### Overview of cell type diversity in the 40% male pupal hindwing

Our analysis of Dsx shows that male wings differentiate two derived scale types with specialized ultrastructures, the UVI and lanceolate scales. Next, we used single-nucleus RNA sequencing (snRNA-seq) to gain further insights into the molecular basis of this diversity of cell types in the male wing. We chose to sequence the hindwings of a single *C. eurytheme* male individual at 40% pupal development, a stage where Bab is consistently repressed in the dorsal cover scales that give rise to the UVI state (*28*). Quality control showed low-contamination of mitochondrial reads following filtering at the 3% threshold (**Fig. S4**), resulting in 2,961 filtered cells. Unsupervised clustering yielded 9 robust clusters, as visualized here in UMAP reduced-dimensionality space (**Fig. 6A)**. We then used *FindMarkers* functions of Seurat version 5 to identify differentially expressed features between these clusters (**Data S1**), and annotate them based on marker gene enrichment and the known function of their orthologs in *Drosophila*. Two minor clusters, dubbed *Misc1* (N = 44 nuclei) and *Misc2* (N = 35 nuclei), remain unannotated and will require further work for confident assignment of cell types within them (see Discussion). The 7 remaining clusters consist of two epithelial cell types dubbed *Wing_epi* and *Trch_epi*, and 5 clusters related to SOP subtypes - namely one socket cluster (*Socket*) and 4 scale sub-types (*Scale1* to *Scale4*), as detailed below. Of note, scale cell types showed higher numbers of mapped reads and detected genes compared to non-scale cell types (**Fig. 6B**), likely due to differences in ploidy levels. Non-scale types also showed a higher level of mitochondrial DNA (mtDNA) contamination relative to scale types (**Fig. 6C**).

**Figure 6.**
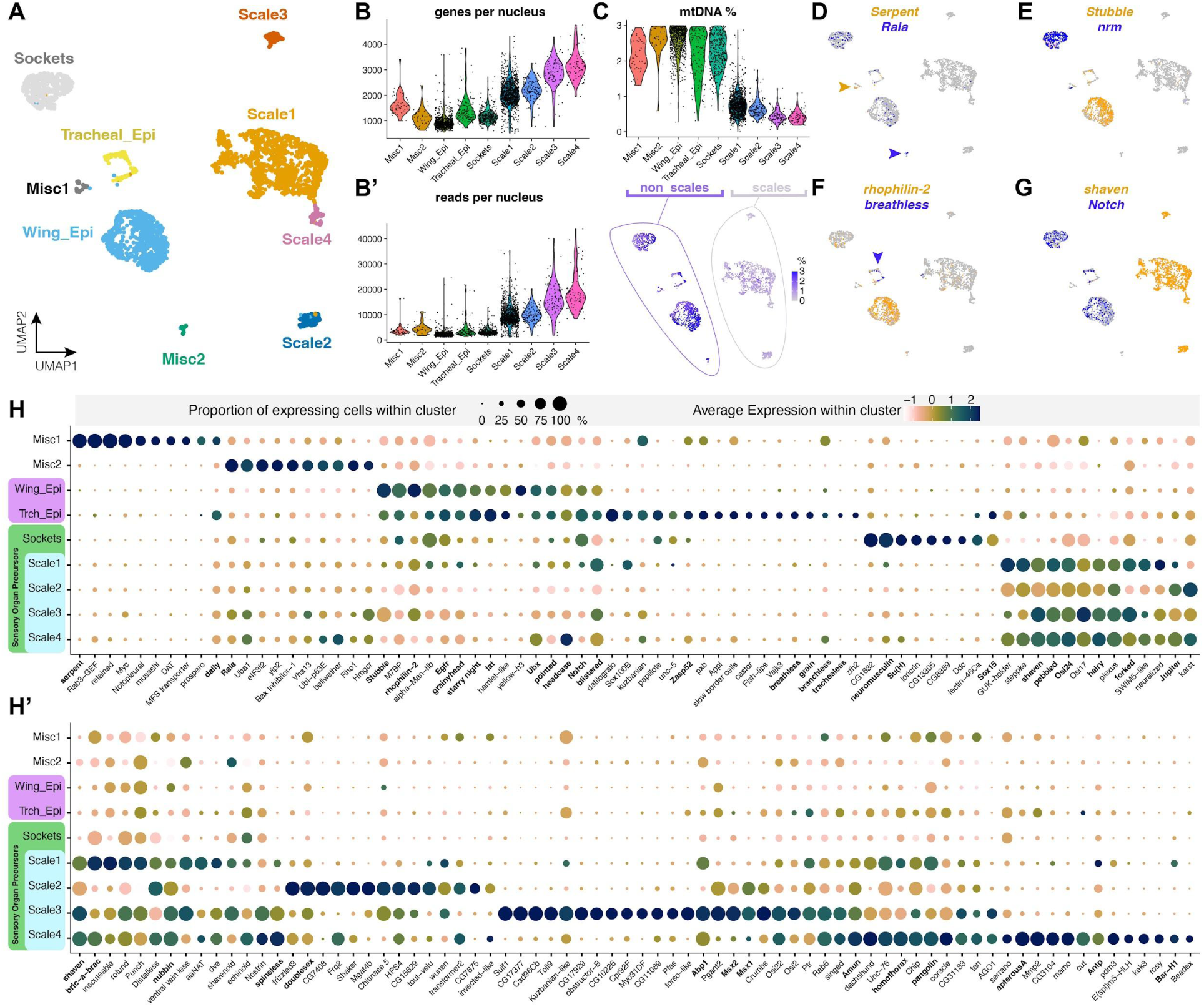
Single-nucleus transcriptomic profiles of major cell types in the *C. eurytheme* male 40% pupal hindwing. **A.** UMAP plot representing the 9 clusters resolved across a total of 2,961 cells. **B-B’.** Violin plots of the number of unique genes (B) and RNA read counts (B’) per nucleus. **C.** Violin and heatmap plots highlighting differences in mtDNA content across non-scale and scale clusters. **D-G.** Heatmap plots showing the expression of key marker genes (bold : see main text for citations). *Serpent* and *Rala* (arrowheads) in two small unannotated clusters, *Stubble* (epithelial cells), *neuromusculin* (*nrm*, socket cells), *rhophilin-2* (wing epithelium), *breathless* (tracheal epithelium), *Notch* (non-scales), and *shaven* (*sv*, scales). **H-H’**. Dot plots featuring the expression profiles of 150 differentially expressed genes, chosen among top markers of cluster or cell type identity. Panel H includes top markers for non-SOP clusters, socket cells, and scale cells ; panel H’ focuses on differentiators between scale sub-types. Dot size reflects the percentage of cells in which gene expression was detected. Color coding allows a relative comparison of gene expression levels within a cluster (horizontal comparisons), but is not proper for vertical comparisons between clusters. Gene names in **bold** : see main text for references.

The bulk of cells providing structural integrity to the wing surface, to the inner surface of the lacunae (luminal tunnels invaded by trachea), and to the tracheal system are all epithelial, diploid cells. Consistent with this, both tracheal (*Trch_epi,* 140 cells) and wing-related (*Wing_epi,* 780 cells) epithelial clusters shared top marker genes that are epithelial cell markers in flies (**Fig. 6**), such as the protease gene *Stubble* involved wing epithelial remodeling (*55*), the EGF signaling pathway genes *pointed* and *EGFR* (*56*), the wing cell polarity factors *fat* and *starry night* (*57*, *58*), and the genes *grainyhead*, *headcase* and *blistered* (*56*, *59–62*). *Ultrabithorax* (*Ubx*) is known as an epithelial marker in butterfly pupal hindwings (*36*) and was enriched in both epithelial clusters. Notch showed an epithelial and socket cell signal consistent with our previous study (*34*). In addition to their shared epithelial gene expression profile, the two clusters differ by the expression of markers specific that are hallmarks of tracheal tip cell growth and tracheal branching, including the tracheal progenitor selector gene *trachealess*, the FGF ligand/receptor pair *branchless/breathless*, the proteoglycan *dally* involved in tracheal FGF signaling, and the cytoskeletal factor *Zasp52* (*63–66*). It is also noteworthy that some *Trch_epi* markers such as *datilografo*, *APP-like* and *slow border cells* were not expected in a trachea-related tissue based on current knowledge, indicating possible evolutionary divergence with *Drosophila* processes. Further investigation will be required to refine the range of cellular identities of the *Wing_epi* and *Trch_epi* clusters, for example to decipher the dynamic complexity of tracheal development, and the differences between wing-membrane and lacunar epithelia in butterflies.

Developmental studies have shown that in butterflies, the SOP lineage has fully differentiated into arranged rows of scale and socket pairs by 30% of pupal development (*37*). This differentiation was well resolved in the 40% stage single-nucleus transcriptome, with a clear cluster of 412 socket cells, based on the expression of the markers *Su(H)*, *Sox15*, and *neuromusculin* (*nrm*), previously associated with *Drosophila* socket specification (*67*, *68*). A total of 1,550 nuclei from scale-building cells were distributed across four distinct clusters that shared the expression of the trichogen master gene *shaven* (*sv*), also variably annotated as *Pax2/5* in insects (*33*, *69*, *70*). Scale-related clusters also shared additional marker genes associated with trichogen development in flies such as *pebbled* (*peb*) (*71*), *Osi24* (*72*), *Jupiter* (*73*), and *Amun* (*74*), supporting the proposed homology between the shaft of mechanosensory bristles with the scale derivative of butterflies and moths (*29*, *31*, *34*). The next section focuses on refining the divergence between groups of scale cell precursors.

### Transcriptome heterogeneity among scale subtypes in the 40% pupal male hindwing

The preliminary differential expression analysis of whole-wing nuclei suggests that scale cell precursors (*sv*^+^/*peb*^+^/*Osi24*^+^) resolve into at least four subtypes, including the large *Scale1* cluster. To delve into the processes of color scale differentiation, we bioinformatically isolated *Scale1-2-3-4*, re-normalized gene expression counts, and reclustered this subset of nuclei to augment the resolution and separation of scale cell subtypes (**Fig. 7A**). All 8 scale clusters expressed canonical scale cell precursor markers such *sv*/*peb*/*Osi24* (**Fig. 7B**). The *Scale1* cluster forms five subclusters numbered *Scale1a* to *1e* for which we have not resolved defined identities at the moment, except for *Scale1d*, which is marked by Antennapedia (Antp) and the pterin repressor Bar-H1 (*22*) and seemingly encompasses the white scales of the wing coupling region (**Fig. 7C-D**). *Scale4* likely corresponds to hairlike scales of the dorsal surface (**Fig. 7D-E**), based on the expression of *homothorax* and *apterous-A* (*75*, *76*), and immunolocalization of Cut (**Fig. 7F-J’**). *Scale1b* expresses *nubbin* and may correspond to a ground scale cell type (**Fig. 7K-L**).

**Figure 7.**
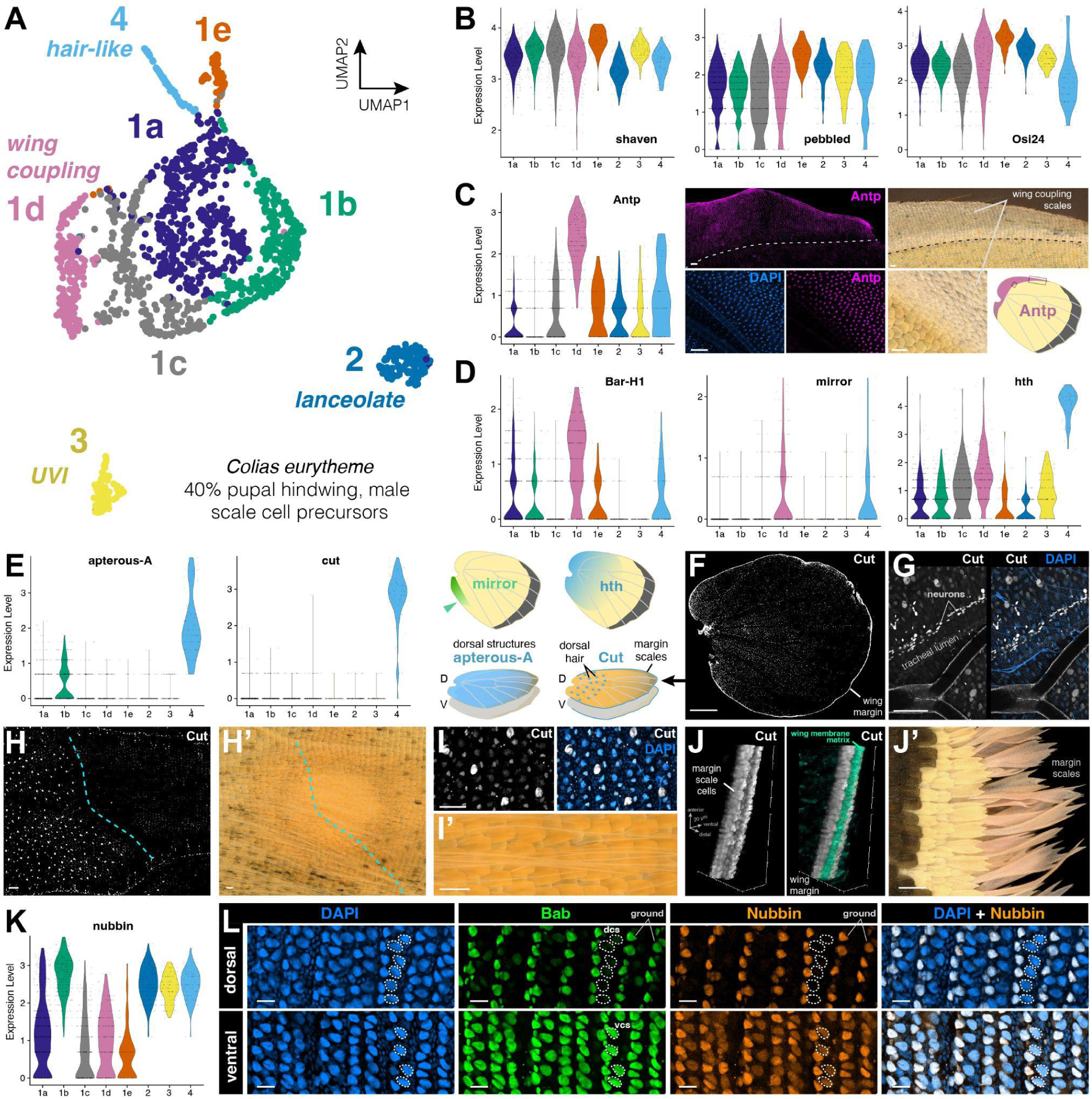
Diverging gene expression profiles of male scale cell precursors at 40% development. **A.** UMAP plot representing the 8 sub-clusters of scale cell precursors across a total of 1,550 cells. **B.** Violin plots for scale markers (*sv*, *peb, Osi24*) and developmental factors showing differential expression patterns across scale subtypes (see text) markers (all scales). **C.** Violin plot of *Antp* expression showing enrichment in *Scale1d*, and Antp immunofluorescence (Antp 4C3 antigen) localization to unpigmented wing coupling scales (*77*). **D.** Violin plots for the unpigmented scales marker Bar-H1 (*22*), far-posterior region marker *mirror* (*78*), and proximal region marker *homothorax* (*76*). **E.** Violin plots for *Scale4* marker genes Cut, previously localized to large wing nuclei and wing margin (*33*, *79*), and *apterous-A*, a marker of dorsal-specific structures (*75*). **F.** Immunofluorescent localization of the Cut 2B10 antigen with bright marking of the wing margin, and nuclear signal intespeced in the wing epithelium and in tracheal lumen. **G.** Cut signal in tracheal lumen likely corresponds to chains of differentiating neurons. **H-I’.** Large nuclei with Cut signal likely correspond for dorsal wing hair, restricted to the proximal side of the discal crossvein (dotted line). **J-J’.** Expression of Cut in wing margin scales likely corresponds to the precursors of elongated margin scales. Cyan: staining of the basal wing membrane (non-specific signal obtained after Dve and immunofluorescence). **K.** Violin plot of *nubbin* expression. **L.** Antigenicity of Nubbin 2D4 revealing repression of Nubbin in dorsal scale cells (dcs) and lower signal in ventral scale cells (vcs). Scale bars: Scale bars: C, G, H, H’, I’, J’ = 100 μm ; F = 1 mm ; I, L = 20 μm.

### UVI and lanceolate scale cells are highly differentiated

To further profile the transcriptomes of the two most divergent clusters, we listed 1,006 genes showing significant differential expression (adjusted *p-value* < 0.05; minimum *log2FC* = 1.25; *min.pct* = 0.25) in comparisons between *Scale2*, *Scale3*, and the remaining scale groupings (**Data S2**). Filtering this list down to 145 most statistically significant genes (adj. *p-value* < 10^-50^) allows a heatmap visualization of the genes that are depleted or enriched in these clusters compared to other scale types (**Fig. 8A-A’**). Remarkably, both clusters *Scale2* and *Scale3* share a low expression of the UVI-state repressor Bab, a repressor of the UVI state that is repressed in UVI cells (*28*). One of the two clusters thus likely corresponds to UVI scales, and we used fluorescent Hybridization Chain Reaction (HCR) mRNA detection of marker genes to resolve their identity. The snRNA-seq signal for *Dsx* indicates it is enriched in the *Scale2* cluster alongside *Arylsulfatase*, a more specific marker for this cell population (**Fig. 8B**). Both genes showed strong transcript signals in the dorsal margin areas where lanceolate scales are located (**Fig. 8C-D**). The *Actin-binding protein 1* (*Abp1*) is a negative marker of *Scale2*, and indeed showed a complementary expression to *Arysulfatase*: while *Abp1* is strongly expressed in the orange/UVI area, this signal decreases in the margin and shows a weak marking of alternating scale cells, as expected from an expression in ground scales (**Fig. 8D**). Meanwhile, the *Scale3* marker gene *Sulfatase1* showed a visible association with the dorsal cover scales of the medial wing area (**Fig. 8E**). Together, these results thus annotate the *Scale2* (*Bab^-^* / *Dsx^+^*) cluster as the population of lanceolate scale cells, and the *Scale 3* (*Bab^-^* / *Dsx^-^*) as the UVI scale cell precursors.

**Figure 8.**
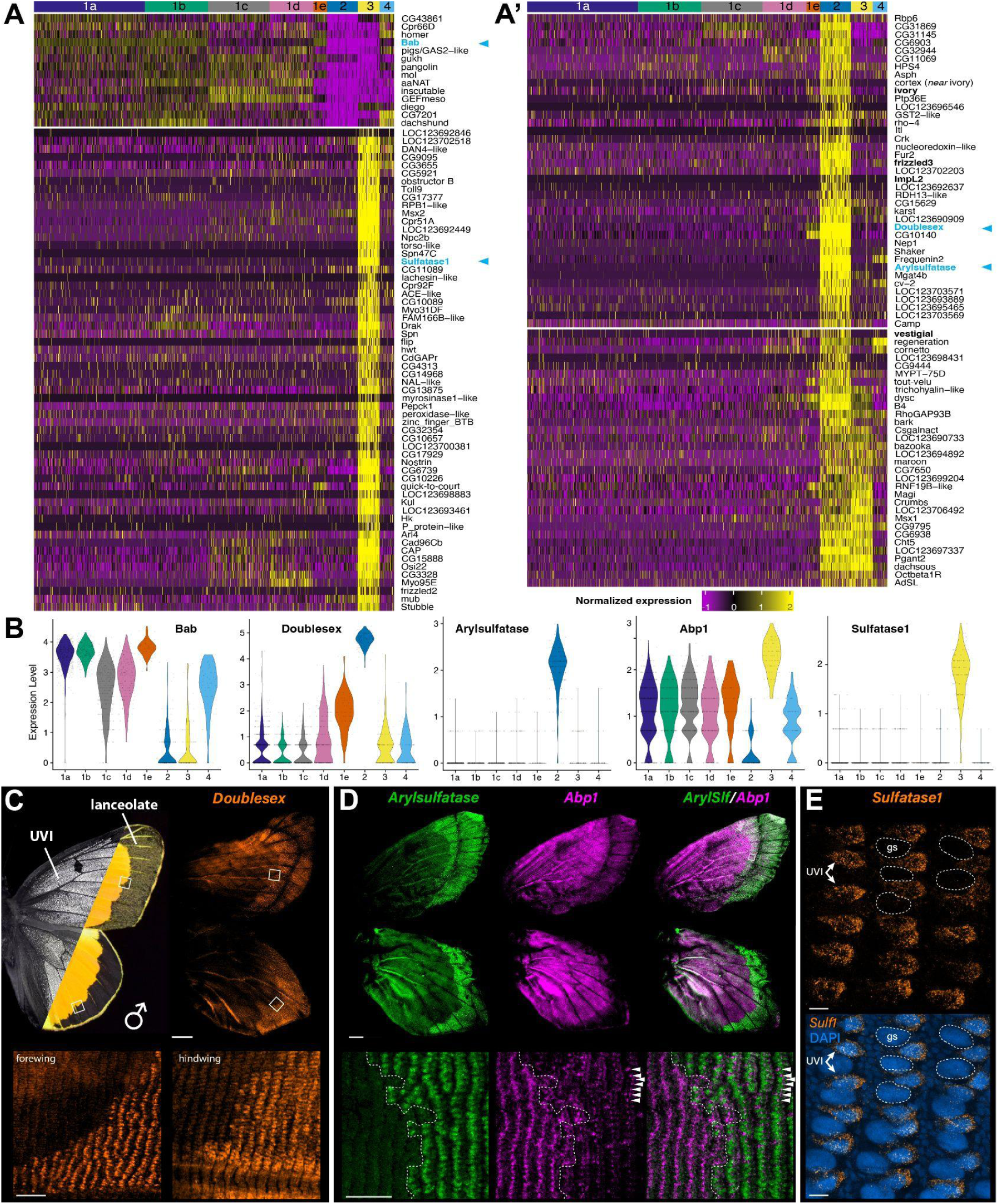
Transcriptional divergence of UVI and lanceolate scale cell precursors. All experiments correspond to wing tissues at 40% pupal development. **A.** Heatmap plot of the 145 most-significant differentially expressed genes in clusters *Scale2* and *Scale3* relative to the remaining scale cell clusters. The top-left panel shows genes downregulated in *Scale2* and *Scale3*. The remaining panels show genes that are enriched in either or both of these clusters. Bold : see text for details. Blue : see panels C-E for spatial expression. **B.** Violin plots for *Bab* and marker genes used for the spatial identification of the *Scale2* and Scale*3* cluster. **C.** HCR localization of *Dsx* mRNA (exons 1-2) in male dorsal wings **D.** HCR localization of *Arylsulfatase* (green) and *Abp1* (magenta), respectively tested as positive and negative markers of the *Scale2* cluster, in male dorsal wings. *Arylsulfatase* is found in the dorsal cover scales of the male marginal region. Arrowheads : ground scale expression of *Abp1* in the marginal region, without overlap with *Arylsufatase.* **E.** HCR localization of *Sulfatase1*, tested as a marker of the *Scale3* cluster, in the medial region of a male dorsal hindwing. Expression is restricted to alternating scale precursor cells corresponding to the presumptive UVI scales, while ground scale precursor cells (gs; dotted lines) are negative. Scale bars: C-D (top) = 1 mm ; C-D insets (bottom) = 100 μm ; E = 10 μm.

In addition, a total of 1,024 genes are differentially expressed between the two *Bab^-^*clusters (**Data S3**), suggesting that while they share the property to be male-specific, UVI and lanceolate scale precursors deploy distinct gene expression programs at the 40% pupal stage. For example, the lanceolate *Scale2* cells are enriched for the *ivory* lncRNA gene and *yellow-c* (**Fig. 8A’**), two potential markers of melanic scales (*80–82*), and express *fz3* and *vestigial*, which mark the periphery of the wing (*83*, *84*). *ImpL2* is the ortholog of *BmIMP*, a gene required for the male-specific splicing of *Dsx* in *Bombyx* (*85*), and is restricted to the *Dsx^+^* lanceolate *Scale2* cells here, suggesting it may play a similar role in butterflies.

### ChIP-seq profiling of Bab genome-wide occupancy identifies potential gene targets

We used ChIP-seq to profile Bab genome-wide occupancy, using fixed nuclei from *C. eurytheme* wings sampled at the 40% and 60% stages, with two biological replicates per stage (**Fig. 9A-B, Data S4**). Following MEME-ChIP motif discovery, the imputed Bab binding matrix recovered the AT-rich sequence profile previously established for *Drosophila* Bab1 using DNAse footprinting assays (*86*), validating the specificity of our ChIP-seq assay. The intersection of predicted Bab occupancy sites at the two stages results in a list of 1,482 candidate genes with at least one stable Bab-binding site in their intragenic interval or immediately adjacent intergenic regions (**Fig. 9C**). Among these, 77 genes are upregulated and 53 are downregulated in the Bab^−^ clusters *Scale2/3* relative to Bab^+^ clusters, thus forming a stringent list of candidate transcriptional targets for Bab (**Data S5**). It is unclear if Bab, a BTB-domain (Broad-complex, Tramtrack, Bric-a-brac) transcription factor, acts solely as a transcriptional repressor, or if it can also act as a co-transcriptional activator (*86–88*). It thus remains uncertain whether the transcriptional targets of Bab should correlate positively or negatively with *Bab* expression, and we illustrate both possibilities with genes that present Bab binding sites and are either enriched or repressed in clusters *Scale2* and *Scale3*.

**Figure 9.**
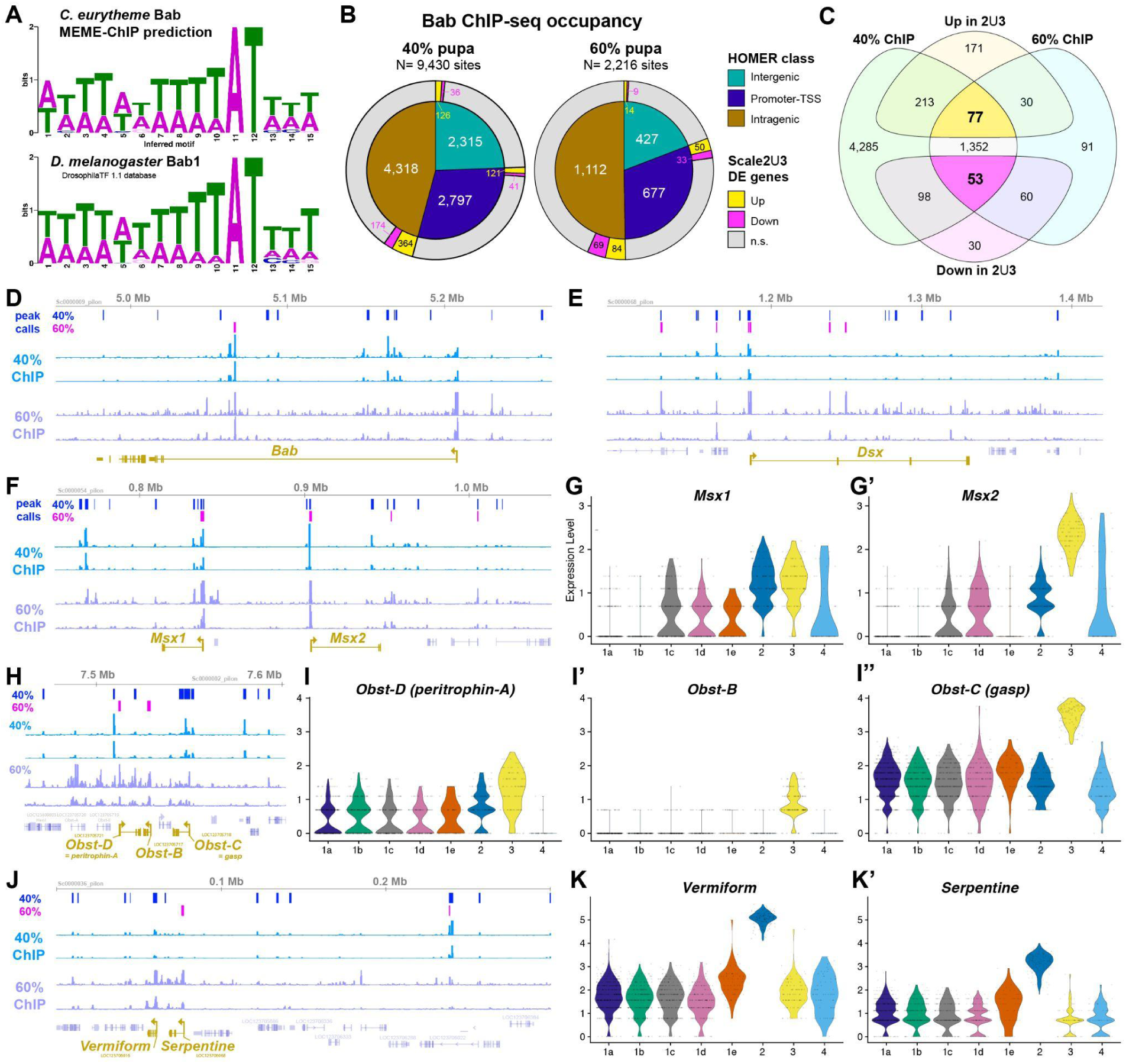
ChIP-seq profiling of Bab identifies potential target genes involved in the differentiation of Bab^−^ scale types. **A.** MEME-ChIP predicted motif for Bab occupancy as inferred from the Bab ChIP pupal wing dataset (top), resembling a previous binding profile for a Bab fly ohnologue (bottom). **B.** Summary of the position of all predicted Bab ChIP-seq binding sites across two datasets, at the 40% and 60% pupal stages. Inner circles : HOMER classification of imputed binding sites relative to gene annotation features. Outer rings : overlap with genes that are among the differentially expressed (DE) genes in *Scale2* (lanceolate) and/or *Scale3* (UVI) relative to other scale types. **C.** Venn diagram featuring the numbers of genes identified as Bab-bound and differentially expressed (DE) in the *Scale2-3* clusters, resulting in an intersection of 77 (up) and 53 (down) DE genes with binding sites identified across two stages. **D, E, F, H, J.** Genomic intervals featuring the Bab ChIP-seq profiles across replicates and peak calls at each stage, including at the promoters of *Scale2-3* markers (bottom, buff color). **G-G’, I-I’’, K-K’.** Violin plot of snRNAseq scale cluster expression for the same marker genes.

For example, the *Bab* locus itself shows multiple Bab ChIP peaks, suggesting it may be auto-regulated by negative and positive feedbacks (**Fig. 9D**). The *Dsx* locus also contains multiple Bab-binding sites (**Fig. 9E**), suggesting that Bab may be an upstream regulator of *Dsx*. Conversely, Dsx is an upstream regulator of *Bab* in flies (*15*, *17*) and also binds a putative intronic enhancer of *Bab* in swallowtail butterflies (*89*). Our data thus suggests that *Dsx* and *Bab* cross-regulate each other, resulting in feedback that may enforce complex dynamics and switches in their expression during sexual differentiation.

The transcription factor genes *Msx1* (*Msh/Drop* ortholog in *D. melanogaster*), and *Msx2* are organized in tandem in butterflies, similarly to *Tribolium* (*90*), and show multiple Bab-binding sites. *Msx2* is a top marker of *Scale3*, while *Msx1* marks both *Bab*^−^ clusters *Scale2-3*, suggesting Bab may act as a repressor at this locus. The conserved family of Obstructor secreted proteins play roles in chitin modifications and cuticle properties (*91–93*). We found that the five members of this family are clustered in tandem in *C. eurytheme*, and that while *Obst-A* and *Obst-E* are not detected in scale cells, *Obst-B*, *Obst-C* (*gasp*) and *Obst-D* (*Peritrophin-A*) are all consistently expressed as markers of the UVI cell cluster *Scale3* (**Fig. 9H-I”**). Strikingly, the promoter regions of these three genes show strong Bab ChIP-seq signals, suggesting Bab may repress the expression of *Obstructor* family genes. Similarly, the chitin deacetylase genes *Vermiform* and *Serpentine* occur in tandem, show evidence of Bab-binding, and are enriched in the lanceolate scale cluster Scale 2 ((**Fig. 9J-K’**).

Overall, these data sketch a broad overview of the potential targets of Bab at both the 40% and 60% stages, leveraging differential expression from both *Bab^-^* cells types – the lanceolate and UVI scale cell precursors. Presently, we do not have functional evidence that low *Bab* expression in *Scale2* cells is required for lanceolate scale specification. In contrast, loss-of-function assays and heterozygous phenotypes of Bab alleles show it is necessary and likely sufficient for UVI scale repression (*28*). We thus sought to further refine the set of potential regulatory targets of Bab by focusing on genes that are differentially expressed in the *Scale3* UVI cells (UVI-DE). Interestingly, genes with Bab ChIP signals are significantly more likely to be UVI-DE (**Fig. 10A**). Stringent filtering criteria result in a list of 87 UVI-DE genes with 3 or more Bab ChIP binding peaks (**Data S6**). Within this set, 17 genes have known functions in *Drosophila* that relate to cytoskeletal dynamics or cuticle formation (**Fig. 10B,C**), two key processes that we expect to regulate ridge spacing and multilayering. For example, the loci encoding the nuclear hormone receptor Eip75b and cytoskeletal regulator Multiple wing hair (Mwh) each show more than 50 Bab ChIP sites, and are upregulated in UVI scale precursors, suggesting that Bab directly acts as a transcriptional repressor at these genes.

**Figure 10.**
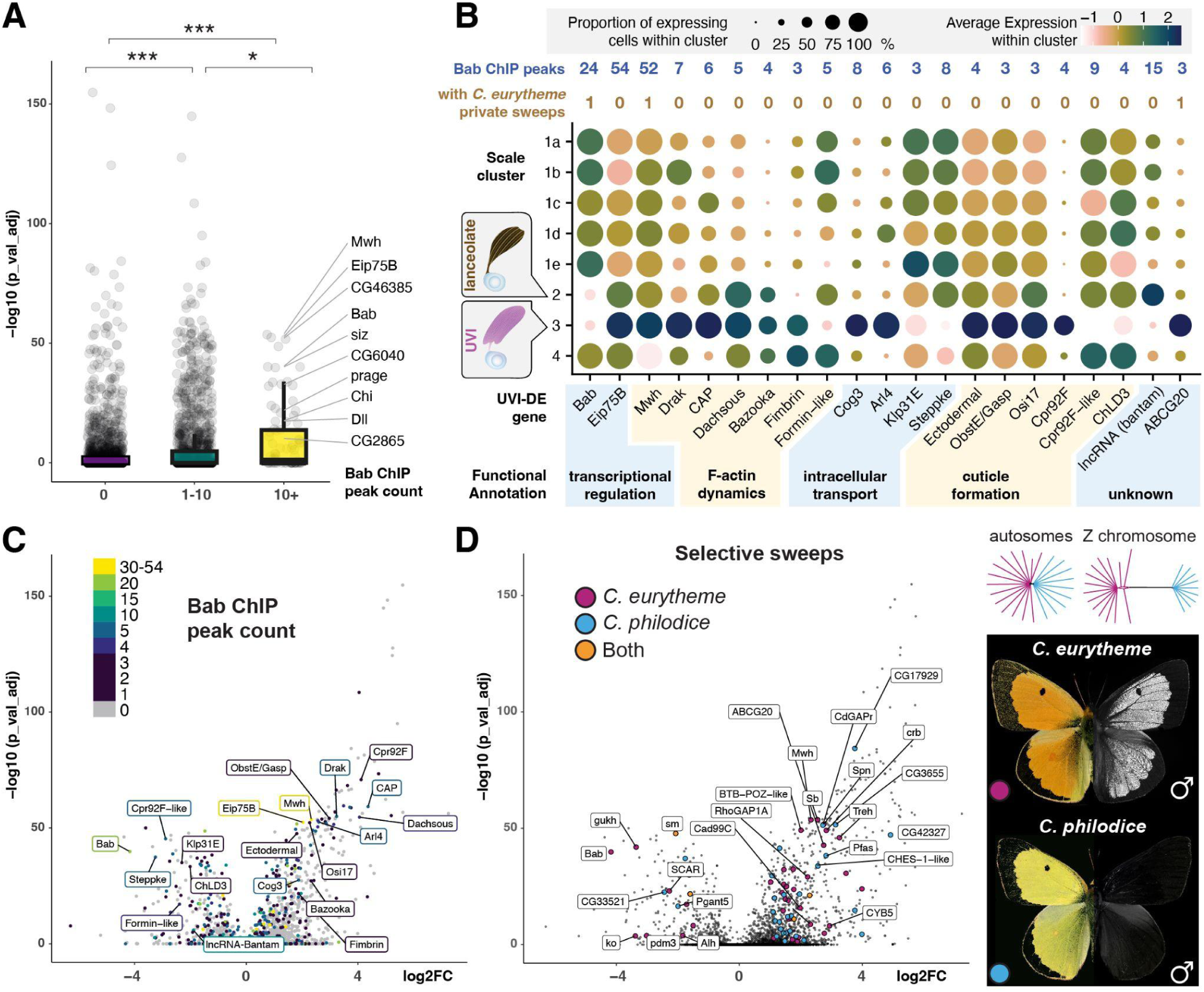
Potential targets of Bab regulation in the UVI scale type. **A.** Box and whiskers plot of probability of differential expression, binned by number of called Bab ChIP sites in proximity to each gene (Wilcoxon rank sum tests; * : *p* < 0.01 ; *** : *p <* 1e-15) **B.** Expression dot plot visualization of selected UVI-DE genes (*log2FC* > 1.8, adjusted *p* < 0.01) with at least 3 Bab ChIP peaks and potential roles in ultrastructural specialization. **C-D.** Volcano plots of UVI-DE genes, colored by number of Bab ChIP sites (**C**), and by the presence of selective sweeps in a sympatric population of *C. eurytheme* x *C. philodice* (**D**). These two morphospecies hybridize and show genome-wide admixture across autosomes (dendrograms reproduced from (*28*)), resulting in a population polymorphic for UV iridescence (present in *C. eurytheme*, absent in *C. philodice*).

### Differentially expressed genes experiencing selective sweeps

*C eurytheme* hybridizes in sympatry with its UV-negative sister species *C philodice* in the Eastern US (*28*). We reasoned that key downstream loci in the formation of UV scales could also show signals of selection that are present in *C. eurytheme* but absent in *C philodice.* Selective sweeps were calculated as the *μ* statistic in RAiSD (*94*) and intervals called with *bedtools*, taking a threshold of the top 1% of *μ* values. 449 sweeps were detected in total, and 63 were found only in *C. eurytheme*. Of 11,647 unique ChIP binding sites, just 29 were found to be within selective sweeps, and of these, 4 are private to *C. eurytheme*, one is private to *C. philodice*, and 24 have undergone a sweep in both species (**Data S7**). Notably *Stubble (Sb),* a determinant of bristle actin bundle number in flies (*95*), has two ChIP peaks that have undergone a sweep in *C. eurytheme* and not *C. philodice*. Most sweeps do not directly intersect ChIP binding sites, but 313 sweeps near 287 genes are in proximity to both ChIP binding sites and selective sweeps, including 58 UVI-DE genes (**Fig. 10D, Data S8**). Notably, *Mwh*, *Sb*, *Bab* and *ABCG20* (**Fig. 10B**) all have private sweeps in *C. eurytheme*, along with *pdm3*, a gene known to be involved in patterning and pigmentation in other butterflies [34], and SCAR, an Arp2/3 interacting protein involved in bristle intracellular patterning in flies (*96*). The combination of differential expression, binding by Bab, and signals of selection on this subset of UV scale-expressed genes provide an initial glance at the candidate genes that fine-tune the specialized ultrastructure of UV scales.

## Discussion

In this study, we tied scale sexual dimorphism to context-dependent functions of Dsx, and identified two male-specific scales types that are demonstrably regulated by the expression of DsxF and Bab (the UVI scales), or by DsxM/F (the lanceolate scales). Then we delved into the single-cell transcriptomics of the male wing tissue and recovered the developmental precursor cell populations of these two derived scale types, providing new insights on how cell-autonomous sexual identity is leading into specialized ultrastructures.

### Multitiered roles of Dsx in wing sexual dimorphism

CRISPR and RNAi phenotypes resulted in concordant phenotypes, and highlight the versatility of Dsx in mediating the sexually dimorphic differentiation of UVI and lanceolate scale types, as well as the patterning of marginal melanic bands that are proper to each sex (**Figs. 5A**, **5F**). The Wnt and Toll pathways, transcription factors Spalt and Distal-less, and the melanic scale determinant *ivory/miR193* have been implicated the patterning of pierid marginal bands (*82*, *97–101*), and future work could explore how Dsx isoforms regulate these processes in early *Colias* pupae.

Lanceolate scales are a male-specific scale type with a derived ultrastructure. As dorsal cover scales specific to the marginal bands of males, this state is positionally reverted to a typical melanic scale phenotype in male *Dsx* perturbation assays, and vice-versa in females. Thus, we conclude that lanceolate scales evolved in unison with both activating input from DsxM and inhibitory effects of DsxF. This dual regulation is consistent with current models suggesting that Dsx targets an identical set of genes between sexes, but where co-factors specific to either isoforms – such as Intersex (IX) for DsxF – confer opposite transcriptional effects on gene expression (*15*, *102*). In future studies of *Colias* wing dimorphism, it will be interesting to combine comparisons of single-nucleus transcriptomes between both male and female tissues, to establish the occupancy profile of Doublesex in each sex (*89*), and to further test the function of potential co-factors such as the lepidopteran ortholog of IX (*103*).

### UV dichromatism is DsxM-independent

Bab is required to prevent UV-iridescence in most scales, and is repressed in the orange dorsal cover scales of male *C. eurytheme*, thus allowing UV-iridescence in a small subset of scales (*28*). Previous work on *Drosophila* sexual dimorphic traits such as the pigmented abdomen, sex combs, and gonads, collectively suggest that DsxF and DsxM have an activating and repressing effect on *Bab*, respectively (*17*, *18*, *104*, *105*). Our experiments rule out a repressing role of DsxM on Bab because Dsx perturbation did not induce loss of UV-iridescence in males. In addition, HCR mRNA detection assays, immunofluorescence, and snRNAseq did not detect Dsx in the UVI cell precursors, ie. the male dorsal cover scales of the medial region. Of note, while we observed a marginal reduction in the UVI field in the vicinity of the peripheral patterns, we extrapolate that this was due to a change in the identity of the marginal patterns rather than a direct effect of DsxM on Bab. The next section thus focuses on the effects of Dsx on cell-autonomous UVI scale repression in females.

### DsxF recruitment enables female-specific repression of UVI patterns

Sexually dichromatic butterflies show a general trend where female wings are usually less colorful than the male ones (*106*), except when colors have an aposematic function. Accordingly, UV iridescence in pierid butterflies is either male-specific (*e.g.* in *C. eurytheme*), or when it is found in females, it is in a much reduced or residual state than in males (*28*, *107*). Here, we found that the medial cover scales express Dsx in females only, where it is required for the activation of Bab, which in turn represses the UVI fate.

In other words, sexual differences in UV-iridescence involve the recruitment of an off-switch in females, rather than a male-specific activation. As noted above, DsxM is a known repressor of Bab in *Drosophila* (*17*), so it appears at first that the recruitment of DsxM to repress Bab in male dorsal cover scales could have led to the evolution of UVI. This turns out to be unlikely, and the mechanism we identified instead makes sense if one considers that the evolution of UVI sexual dichromatism occurred in two steps. In this scenario, UV-iridescence first originates in both sexes, and involves a loss of Bab expression in the dorsal cover scales of the medial region. Later, reduction of UVI patches evolves in females by recruiting Dsx in the dorsal cover scales, re-activating the expression of Bab. This off-switch mechanism went a further step in *Zerene cesonia*, a close relative of *Colias*, where a duplicated copy of *Dsx* evolved a truncated female-specific transcript that is highly expressed in the female dorsal scales, in a pattern consistent with UVI-repression (*108*). We thus postulate that Dsx did not contribute to the origin of the UV scale type itself, but that its female-specific products were recruited to reduce the size of the UVI patch in females. Crucially, this model is consistent with phylogenetic models that place the origin of UV iridescence in an early ancestor of Pieridae, followed by secondary reduction of UVI patterns (*20*).

How does this mechanism fit more broadly into evolutionary thinking? In a decade-long debate about the causes of sexual dichromatism (primarily in birds and butterflies), Arthur Wallace invoked the importance of natural selection to decrease conspicuous traits in females, while Charles Darwin favored the supremacy of sexual selection (*109*). Because males and females are often subject to different selective pressures, it is commonly accepted that the two explanations are not mutually exclusive (*110*, *111*). In our evolutionary model, sexual selection remains a key force to explain the origin of UV-iridescence and its persistence in males (*24*, *112*), as defended by Darwin, while the spatial reduction of this trait in females supports a role for natural selection as championed by Wallace. For example, as UV iridescence may make butterflies more visible to passerine birds with ultraviolet-vision (*113*), it is possible that its reduction in females is linked to predator avoidance. The recruitment of DsxF in UVI scale repression provides a mechanism for this sex-specific reduction.

### Using snRNAseq to explore wing cell type diversity

The enormous diversity of butterfly wing patterns largely amounts to the ability of these organisms to differentiate various scale types that vary in coloration, and to organize them in space during development. Scales emerge from a single scale cell precursor during development, and we expect that single-cell approaches to scale differentiation will uncover new insights on how tissues organize complex spatial patterns during development and how new traits evolve. In this study, single-nucleus transcriptomics allowed the detailed profiling of nearly 3,000 cells, about half of which were scale cell precursors with low mtDNA contamination and high read counts. Each snRNAseq sample consisted of a single hindwing dissected from a male *C. eurytheme* pupa, and the pooling of several individuals is unnecessary with this technique and tissue. Subsetting the initial clustering to select for scale clusters, characterized by *sv* expression at this stage, enables the detailed analysis of scale cell precursor transcriptomes in ways that have not been possible with bulk RNAseq methods, leading to the mining of top marker genes from well differentiated clusters and generated new hypotheses regarding cell type specification and specialization.

### Transcriptomics insight on the differentiation of specialized scale ultrastructures

We found that two scale clusters lack *Bab* expression, and correspond to male-specific, Dsx-dependent scale types – the UVI and lanceolate scales. Each of these types show ultrastructures that diverge from canonical scale types, with extreme variations in the spacing of longitudinal ridges (**Fig. 1D**), and the remarkable multilayering of the UVI scale upper lamellae (**Fig. 1J**) that yields their structural coloration (*41*, *114*).

Previous studies have shown that condensed actin filaments pre-pattern the density of ridges in butterfly scales, and then template the formation of iridescent elaborations (*115–118*). Genes that are differentially expressed in the UVI scales provide a wealth of candidate modulators of actin dynamics including *shavenoid*, *Abp1*, *singed* (**Fig. 6H’**), which are known to modulate actin assembly and ridge patterning in *Drosophila* bristles (*96*, *119*, *120*). We found strong evidence of Bab occupancy at the UVI scale marker genes *fimbrin* and *mwh,* two key determinants of actin bundling and bristle morphology in flies, indicating they are direct targets of Bab repression (**Fig. 10B**). Similarly, we suggest that Bab-bound DE genes with annotated functions in vesicular trafficking and cuticle formation – such as *Osi17*, *Cpr92F*, and the chitin deacetylase *ChLD3* (*121–123*) – may be direct targets of Bab regulation that participate in the sculpting of the nanostructures on the scale upper surface, including the stacking of UVI ridge lamellae. This is further strengthened by the fact that a number of these genes also show evidence of selection specifically in *C. eurytheme* and not its UV-negative sister species *C. philodice* (**Fig. 10D**), implying either that regulatory connections are being maintained by purifying selection or that positive selection for network interactions permissive for UVI scale specialization are under positive selection. While the function of these potential effectors will require further investigation, the application of snRNAseq to mid-pupal stages of butterfly wing development promises to be an exciting avenue of research for understanding how complex exoskeletal ultrastructures form and diversify.

## Methods

### Butterflies

*C. eurytheme* females were collected from an organic alfalfa field (Buckeystown, MD), and left in small cylinder cages with alfalfa for oviposition. Eggs were collected manually, washed 1’ with Benzalkonium Chloride 5%, rinsed and dried, left to hatch at 25°C, 60% humidity, 14:10 h day:light cycle and fed with Black Cutworm artificial diet (Frontier Agricultural Sciences) supplemented with dry alfalfa powder, before transfer to Lana woollypod vetch (*Vicia villosa*) in a greenhouse environment (*23*). Fifth-instar wandering larvae were moved to a growth chamber set at 28°C, 60% humidity, 16:8 h day:light cycle, and their time of pupation recorded.

Developmental times are reported as percentages, relative to a mean development time 138 h from pupation to adult emergence at 28°C, or 170 h at 25°C.

### CRISPR mosaic knock-outs

Eggs were collected on alfalfa shoots and leaves, washed, dried, and placed upward on thin strips of double-sided tape, on the inner side of a 1.25 oz. cup lid, and micro-injected following previously reported procedures (*23*, *28*) with an equimolar mixture of Cas9-2xNLS:sgRNA at 500:250 ng/μL. Embryos were left to hatch in cups containing vetch sprouts, and transferred to vetch sprout mats in a greenhouse environment for larval growth.

### RNAi electroporation

Dicer-substrate siRNAs (DsiRNAs) were designed against target gene exons (**Table S2**), ordered at the 2 nmol scales as Custom DsiRNAs with standard purification (IDT DNA Technologies), resuspended at 100 μM in 1x *Bombyx* injection buffer (pH 7.2, 0.5 mM NaH_2_PO_4_, 0.5 mM Na_2_HPO_4_, 5 mM KCl), and stored as frozen aliquots at −70°C until use. Electroporation procedures followed a previously described procedure (*84*, *124*). To target the dorsal surface of male wing pupae, the negative electrode was placed in contact with the droplet of saline positioned on the ventral side of the peeled wing, while the positive electrode was in contact with the agarose pad on the dorsal side of the wing, before electroporation with 5 square pulses of 280 ms at 8 V, separated by 100 ms intervals.

### Genome annotation

The genome annotation available for *Colias croceus* (NCBI:PRJEB42949), was mapped to the best available *C. eurytheme* assembly (NCBI:GCA_907164685.1) using *Liftoff* (*21*, *125*, *126*). A total of 12,908 genes had annotations transferred from *C. crocea* and 7,418 genes were obtained on reciprocal Blast against Flybase protIDs using *-blastx* with *-evalue 1e-50*, retaining best hits against subjects.

### Live nuclei preparation for single-nucleus transcriptomics

We isolated live nuclei using dounce homogenizers following previously published protocols (*34*, *127*). Briefly, two hindwings were dissected from a single male individual (53 h old pupa at 28°C, 40% stage) in cold 1x Phosphate Buffer Saline (PBS) and immediately transferred for sequential douncing in homogenization buffer (250 mM sucrose, 10 μM Tris pH 8.0, 25 mM KCl, 5 mM MgCl_2_, 0.1% Triton-X 100, 0.2 U/μL RNasin Plus, 1x Protease Inhibitor, 0.1 mM Dithiothreitol). The homogenized tissue was transferred to a new 1.5 mL tube and spun down in a fixed-rotor centrifuge at 4°C for 10 min at 1,000 *g*, before resuspending in 500 μL nuclei suspension buffer (PBS, 1% Bovine Serum Albumin, 0.2 U/μL RNasin Plus). The nuclei suspension was filtered using a 40-μm PluriSelect filter and 10 μL were drawn for staining with Trypan Blue. Nuclei quality was assessed based on membrane integrity and nuclei were counted using a hemocytometer, yielding a concentration within the target range of 600-900 nuclei/μL. Nuclei suspension was processed for cDNA library preparation with a target output of 3,000 nuclei per sample.

### snRNAseq library preparation and sequencing

The droplet-based 10X Genomics Chromium Next GEM Single Cell 3ʹ Reagent Kit v3.1 with Dual Indexes was used to prepare multiplexed cDNA libraries from nuclei suspensions (*34*). Libraries were pooled for PE150 sequencing on a Novaseq S2 flow cell at a target depth of 200,000 reads/nucleus. Raw reads are accessible at the NCBI SRA (NCBI:SRS12272185).

### SnRNAseq data preprocessing and analysis

*BCL* files were demultiplexed into *fastq* files using *bcl2fastq*. Raw and filtered CellRanger-generated count matrices with the flag *--include-introns* were used for downstream analysis and are available in an online data repository (*128*). Seurat v5.0 was used for preprocessing of Seurat objects, first filtered to remove low-quality Gel Beads-in-emulsion (GEMs) with *nCounts* < 300, *nFeatures* < 3,000 and *percent.mt* > 3%. Library sizes, number of per-cell features and proportion of reads mapped to mitochondrial genome were used as quality control metrics (Islam et al. 2014; Ilicic et al. 2016). The sample at 40% pupal development was normalized using *SCTransform* with an additional regression of nuclei based on percentage of mitochondrial reads (*vars.to.regress* = “*percent.mt”*), before performing *RunPCA* and *RunUMAP* using top 20 PCs, after which *FindNeighbors* and *FindClusters* functions were invoked to generate 2D layouts of graph-based clusters. Clustering resolution of 0.1 was used after examining cluster stability using *R/clustree* on clustering resolutions at 0.1, 0.2, 0.5, 0.7, and 1.0.

For cluster annotation of recovered nuclei from the 40% sample, we identified differentially expressed genes using *FindAllMarkers* in Seurat (Wilcoxon Rank Sum test with Bonferroni correction for multiple testing; adjusted *p* < 0.05). Marker genes were identified based on genes detected in a minimum of 25% of the cells within the cluster and a *log2FC* > 0.25 between the cells in the cluster and all remaining cells. Focusing on scale-building nuclei, the whole object was subset for *shaven-*expressing nuclei. Preprocessing steps were performed as previously mentioned but using 0.2 resolution for *FindClusters*. Heatmap was generated using *R/ComplexHeatmap* (*129*, *130*) and genes were ordered via unsupervised hierarchical clustering of 4 k-means groups.

Following cluster identification, we subset the whole object for scale clusters *Scale1-4* using expression of canonical markers *sv* and *ss*. Normalization with *SCTransform* was performed on the original, filtered count matrix as mentioned previously, following *RunPCA* and *RunUMAP* with top 10 PCs, before *FindNeighbors* and *FindClusters* were performed with a clustering resolution of 0.1. *FindMarkers* comparing each cluster of interest with the rest of the subsetted object adjusted *p* < 0.05, *log2FC* > 1.25, *min.pct* = 0.25) was used to identify marker genes.

### Preprocessing and analysis of Chromatin-Immunoprecipitation (ChIP)-sequenced data

ChIP-seq libraries were prepared from live pupal wings (**Supplementary Text**), and sequenced as PE42 reads for samples at 40% and PE37 reads for samples at 60%, available on the NCBI SRA, PRJNA1148116). The *C. eurytheme* genome was indexed using *bowtie2* (2.5.3). *FastQC* was used to assess the quality of fastq files. Trimmed reads were aligned to the genome using *bowtie2* (2.5.3), sorted using *samtools* (1.15.1), filtered, and *MarkDuplicates* using *picard tools* (2.26.8) from GATK (4.2.4.0) was invoked to remove duplicate reads. Filtered and uniquely mapping reads were used for peak calling using MACS3, with options *q = 0.01*, *BAMPE*, and an effective genome size of 328,651,476 bp. Relaxed peak calling with a *p = 0.05* threshold was performed, followed by further statistical correction using the Irreproducible Discovery Rate framework (version 2.0.4.2), which assessed the reproducibility of called peaks between the two replicates based on a fraction of false positive peaks (*131*, *132*). This generated a final list of peaks for each timepoint at the maximum False Discovery Rate (FDR) of 0.05 (**Data S4**). Peak annotation was performed using HOMERv4 *annotatePeaks.pl*, to assign each peak to a genomic region of a nearby gene (*133*). For visualization in *IGV*, the *deeptools* (3.5.4) function *bamCompare* was used to subtract input *bam* files from each corresponding replicate and derive *bigWig* files for each sample (*134*). To identify conserved motifs within peaks across four samples, *fasta* files for peak intervals were extracted from *IDR* output using *bedtools* (2.28.0) function *getfasta*, and served as the input for motif discovery using the Combined Fly reference set in MEMECHIP (*135*). Finally, gene lists from **Data S2** and **S4** were merged to identify candidate genes regulated by Bab, with differential expression in the *Bab^-^* scale clusters and Bab-binding ChIP peaks present in both pupal stages, 40% and 60% development.

### Immunohistochemistry and confocal imaging

Antibody stainings were performed as previously described (*28*). In short, pupal wings were dissected in cold PBS and fixed in fixative (4% methanol-free paraformaldehyde diluted in PBS, 2 mM egtazic acid) at room temperature for 13-20 min, washed four times in PT (PBS, 0.1% Triton-X 100), blocked in PT-BSA (PT, 0.5% Bovine Serum Albumin), incubated overnight at 4°C with primary antibody dilutions in PT-BSA, washed in PT, incubated 2 h with secondary antibody dilutions in PT-BSA (1:500 dilutions, following two short centrifugation steps to pellet down particules before incubation with the wings), washed in PT, incubated in 50% glycerol with with 1 μg/mL DAPI (4′,6-diamidino-2-phenylindole), mounted on glass slides with 70% glycerol or SlowFade Gold mountant under a #1.5 thickness coverslip, and sealed with nail varnish before confocal imaging. Antibodies used in this study were a polyclonal, affinity purified anti-*Colias* Bab (*28*) rabbit antibody (1:100 dilution); several monoclonal mouse antisera (1:50-1:100 ; Developmental Studies Hybridoma Bank) targeting DsxDBD (*54*), Nubbin (*136*), Cut (*137*), and Antp (*138*) ; and a polyclonal, affinity purified anti-Dve (*139*) guinea pig antibody (1:400 ; kind gift of Mike Perry). Secondary antibodies included conjugated AlexaFluor488 anti-Mouse IgG (Life Technologies, CA) at 1:500 dilution, conjugated AlexaFluor647 anti-Rabbit IgG (Life Technologies, CA) at 1:500 dilution, and conjugated AlexaFluor555 goat anti-guinea pig IgG (Abcam, UK). Stacked acquisitions were obtained on a Olympus FV1200 confocal microscope mounted with PLANAPO 20x and 60x objectives. Whole-wing fluorescent microscopy images were obtained with a Zeiss Cell Observer Spinning Disk confocal microscope mounted with a 10x objective (Plan-Apochromat, 0.45 NA) for HCR mRNA stainings, and with an an Olympus BX53 epifluorescent microscope mounted with an UPLFLN 4x objective for immunofluorescent stains.

### Detection of selective sweeps

An admixed population of *C. eurytheme* and *C. philodice* (Buckeystown, MD) was previously resequenced, genotype-called, filtered and analysed (*28*). RAiSD was used to calculate the µ statistic separately on each of *C. eurytheme* and *C. philodice* with default parameters (*94*). Swept intervals were identified by taking the top 1% of µ for each species and merging the called sites with *bedtools merge*. Each called interval was then connected to its nearest gene using *bedtools closest*, and intersected with called ChIP sites with *bedtools intersect*.

## Acknowledgements

We thank Rachel Canalichio and the staff of the Harlan W. Hilbur Greenhouse at the GWU for providing butterfly host plants, Patricia Hernandez and Aleksandar Jeremic for providing access to confocal microscopes, the Duke University Sequencing and Genomic Technologies Shared Resource for assistance with library sequencing, Alejandro Berrio and Carlos Arias for bioinformatics assistance, Hedgeapple Farms for providing access to field collection sites, and Brian Counterman, Vincent Ficarrotta, and Adam Porter for stimulating discussions on ultraviolet dichromatism. **Funding.** This work was funded by the NSF awards IOS-2110532, IOS-2110533, IOS-2110534 to WOM, GAW and AM; the NSF award IOS-2128164 to RDR; the Wilbur V. Harlan Research Fellowship to MT, LSL, and AC; the Smithsonian Institution (SI) Postdoctoral Fellowship in Biodiversity Genomics to JJH.

## Supplementary Materials

### Supplementary Text

#### Data accessibility and remarks on annotated gene names and marker detection

We provide all necessary data for further exploration of the *Colias* single-nucleus transcriptome in an online repository accompanying this article (*128*). This includes the reference genome annotation, inferred gene names, and computed tables for various differential expression analyses. We wish to highlight two technical limitations in our current analysis workflow that can impact future analyses of these datasets.

Gene name annotations do not always indicate direct orthology with traditional model organisms. In our annotation table, we provide two name categories. The first one is based on the best BLASTP hit against the *Drosophila melanogaster* reference protein set. The second one is based on NCBI-provided annotations, derived from the *Colias croceus* annotation. These gene names are based on homology with more phylogenetically distant mammalian cognates. When featuring gene names in our figures, we manually verified reasonable orthology relationships by performing additional BLAST analyses across insects, and favored *Drosophila* gene names when we could identify a one-to-one orthology. Gene names in the provided tables require more caution on a case by case basis, and an insect-wide gene orthology and nomenclature will be required in the future to address these issues.

Lastly, we followed common recommendations to include intronic reads in the definition of transcribed units for our single-nucleus experiment (*140–142*). While this allows the detection of transcripts expressed at low levels and is essential for this type of analysis, this means that overlapping gene annotations can create erroneous allocations of a given signal to a gene name in our DE tables. For example, if *Gene A* is differentially expressed but is nested within *Gene B*, *Gene B* may be reported as the DE gene. With this caveat in mind, it will be important to manually examine the genomic context of candidate genes before proceeding to gene-specific assays, and the definitive resolution of this issue will require innovation in RNA mapping pipelines for this type of analysis.

#### Supplementary methods : PCR genotyping

The chromosomal sex of *Dsx* crispants was determined by genotyping the presence of one or two Z-linked alleles by Sanger sequences. Two Z-linked PCR products (*M13F-fwd1*: 5’-TGTAAAACGACGGCCAGTCTCCGGGATGACTACTTGAC-3’ ; *rev1* : 5’-TGATCTCCGGAGCCATGAAG-3’; *M13F-fwd*2: 5’-TGTAAAACGACGGCCAGTTTAACACACGACTGAGACCC-3’ ; *rev2* : 5’-TCGGTGTGAGGCTCAGGTAC-3’) were amplified from a single leg using Phire Tissue Direct PCR Master Mix, column-purified, and sequenced using the M13F universal primer. Sequences with double-chromatograms were inferred as originating from ZZ males, while single chromatograms were inferred as females. This method accurately predicted the sex of control individuals of known sex, and matched the inference made from genital morphology in crispant individuals. Genital morphologies did not show asymmetries or visible phenotypes among surviving adults, a negative result that we attribute to possible lethal effects of *Dsx* CRISPR mKOs during development (**Table S1**), as previously observed in other butterflies (*9*, *143*). The presence of indel mutations at the CRISPR targets sites was verified using chromatogram deconvolution with ICE (*144*), on amplicons generated by direct PCR as described above and sequenced with the reverse primer (*Ce_Dsx_sgRNA1*, *fwd* : 5’-GCAATGCTTTTTGCTTGCCA-3’, *rev* : 5’-TTTGGTGTTGTCAGGTACGG-3’; *Ce_Dsx_sgRNA2, fwd* : 5’-GTGGGCTTGTGAAATTGCAT-3’, *rev* : 5’-GCGTGGGGTCATCCAAAAAT-3’).

#### Supplementary methods : Preparation of Chromatin-Immunoprecipitation (ChIP)-sequencing libraries.

A total of 16 pupal wings were dissected from two females and two males at 40% pupal development for 2 library replicates, and similarly 16 pupal wings from 4 individuals at 60% pupal development for 2 library replicates. Tissues from each individual were fixed in 1% formaldehyde in PBS, before adding PBS with 0.135 M glycine for 5 min, then washing with ice-cold PBS twice. All liquid was removed prior to flash-freezing tissues in liquid nitrogen for 30 s. We performed sonication and library preparation for ChIP-sequencing following previous recommendations (*145*). Fixed tissues were dissociated in a sucrose buffer supplemented with Protease Inhibitor (PI) using a dounce homogenizer. Homogenized tissues were then spun at 2,000 *g* for 5 min, and the cell pellet was treated with 1 mL freshly prepared ATAC lysis buffer and PI for 5 min. During this treatment, the cell suspension was pipetted to avoid clumping. Cells in the lysis buffer were spun at 2,000 *g* for five minutes. The lysis buffer was removed, and the pellet was resuspended with 1,000 mL of 1x ChIP Dilution Buffer (Cell Signaling). 150-200 μL of the cell suspension was then distributed into Diagenode 1.5 mL TPX microTubes for sonication. Cell suspensions were then sonicated in < 4°C in a Diagenode Bioruptor UCD-200 (3 x 5 min, 30 s on with *high* setting, 30 s off). Sonicated cell suspensions were then centrifuged at 12,000 *g* to release chromatin, then supernatants from individual tubes were pooled for 20-50 μg of DNA per 1 mL. Sheared DNA was then incubated overnight with 7 μL of 0.550 mg/mL anti-Bab antibody at 4°C, or left untreated for input control samples. Both control and Bab-bound sheared DNA were then treated with 10 μL of Dynabeads Protein A and 10 μL of Protein G (ThermoFisher Scientific Inc. 10001D, 10003D) for a minimum of 2 hours nutating at 4°C. A magnetic rack was used to separate the immunoprecipitated DNA on Dynabeads, washing three times with 1,000 μL of 1x ChIP dilution buffer and twice with 1,000 μL of 1x ChIP Dilution Buffer supplemented with 70 μL of 5M NaCl. Immunoprecipitated DNA on magnetic beads were then resuspended in 1x 150 μL of ChIP Elution Buffer (Cell Signaling Technology) and incubated for 1.5 h at 65°C vortexing every 10 min. After 1.5 h, the supernatant containing the immunoprecipitated DNA was treated with 2 μL of 5M NaCl and 10 μL ProteinaseK, then incubated for 2.5 h to de-crosslink. The samples were then purified using the SimpleChIP Chromatin IP kit (Cell Signaling Technology). DNA libraries were made by normalizing the input to the ChIP-ed DNA, using a NEBNext Ultra Prep II DNA library kit without size selection and with 13 cycles of PCR amplification. Prepared libraries were cleaned with 0.9x Ampure beads and checked for fragment distribution, before sequencing by the Biotechnology Resource Center (BRC) Genomics Facility (RRID:SCR_021727) at the Cornell Institute of Biotechnology on their Illumina NextSeq 500/550 platform. ChIP-seq libraries were sequenced as PE42 and PE37 reads for samples at 40% and 60% pupal development, respectively, and are available on the NCBI SRA (Bioproject PRJNA1148116).

#### Supplementary methods : Adult wing imaging

Adult wings were imaged in the visible range using a Nikon D5300 camera mounted with a 105mm f/2.8D AF Micro Nikkor lens, and a VHX-5000 microscope mounted with VH-Z00T and VH-Z100T lenses. For UV-photography, full-spectrum converted Panasonic G3 camera was mounted with UV-transmitting lenses (*20*, *28*) and used to image wings under the UV-illumination of GE Blacklights 13-Watt T3 Spiral Light Bulbs and 365nm LED torchlights.

---

**Figure S1.**
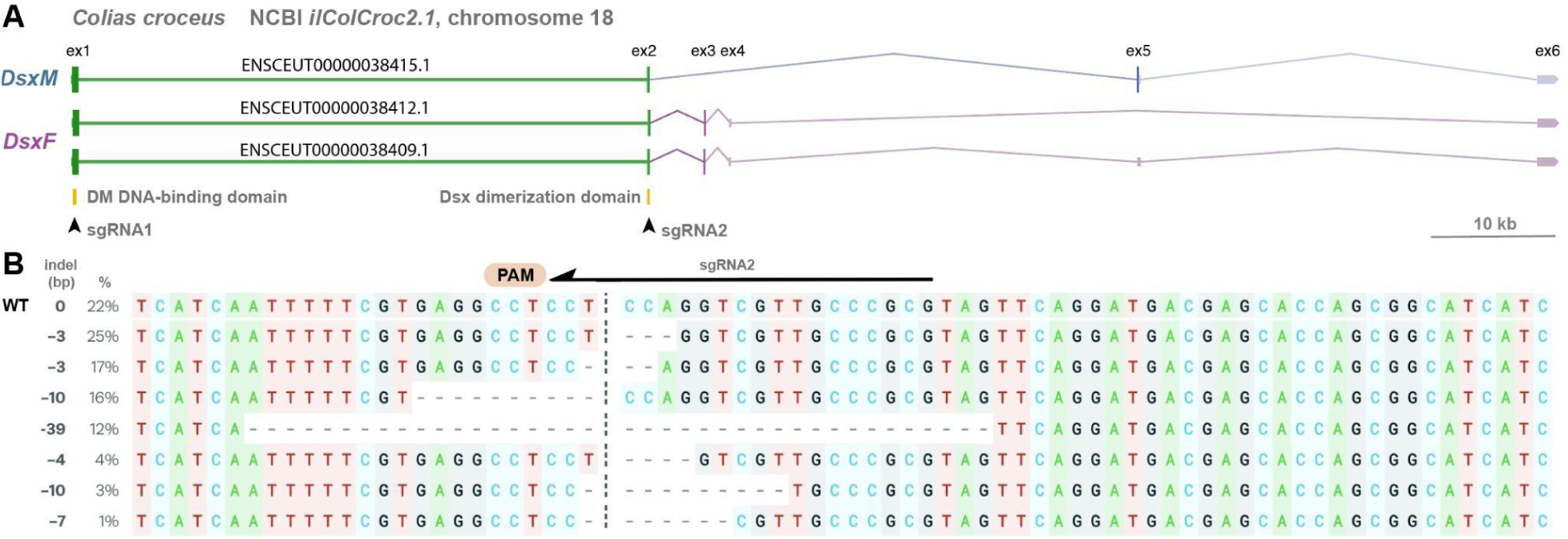
Targeted mutagenesis of *Dsx* monomorphic exons in *C. eurytheme.* **A.** Overview of the *Dsx* locus in the *Colias croceus* genome annotation (*125*). The gene structure is inferred from RNAseq intron-spanning reads available on the NCBI Genome Browser, and features three major isoforms. The male isoform (*DsxM*, ENSCEUT00000038409.1) spans an open-reading frame on exons 1, 2 and 5. The female isoforms (*DsxF*, ENSCEUT00000038412.1 and ENSCEUT00000038415.1) both span an open-reading frame on exons 1, 2 and 3. CRISPR sgRNA targets were designed on the matching version of the *C. eurytheme* genome (arrowheads) and predicted to impact all isoforms. The *exon 1* target overlaps with the region encoding the DM DNA binding domain of Dsx, while the *exon 2* targets corresponds to the Dsx dimerization domain. **B.** Genotyping of a mosaic crispant following the targeting of *Dsx exon2* using Synthego ICE chromatogram deconvolution. Dotted line : predicted cut site.

**Figure S2.**
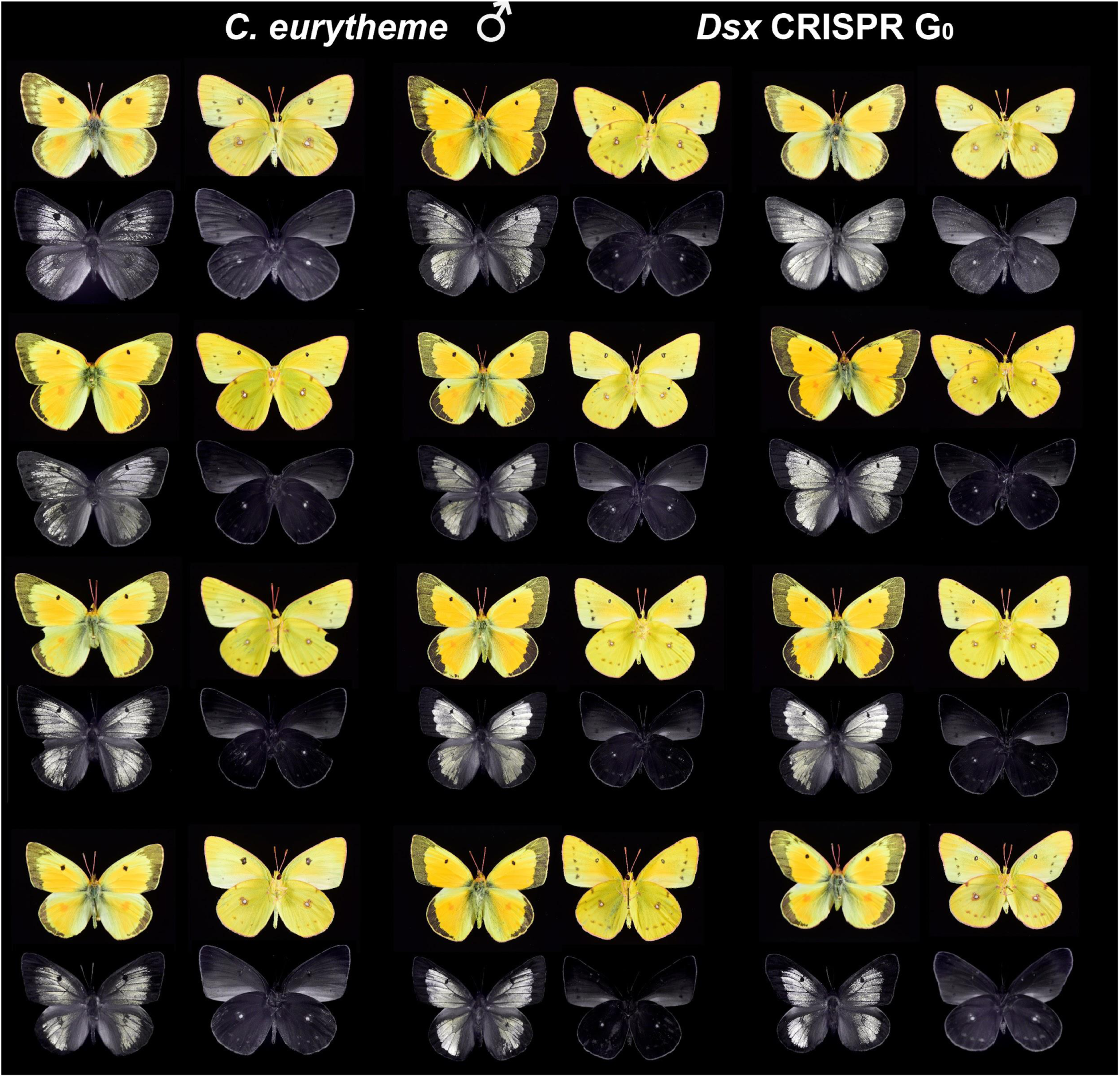
*Dsx* crispant phenotypes in *C. eurytheme* males. *Dsx* mosaic knock-out effects are exclusively visible in the marginal section of the male dorsal wings (left panels), with a proximal extension of the melanic band, a proximal regression of the distal border of the UV iridescent region (also visible in the visible spectrum as a yellow extension), and a transformation of scent-related marginal scales into regular melanic scales (not visible here). There were no visible effects on ventral sides (right panels). Bottom rows : UV-spectrum photography (320-400 nm).

**Figure S3.**
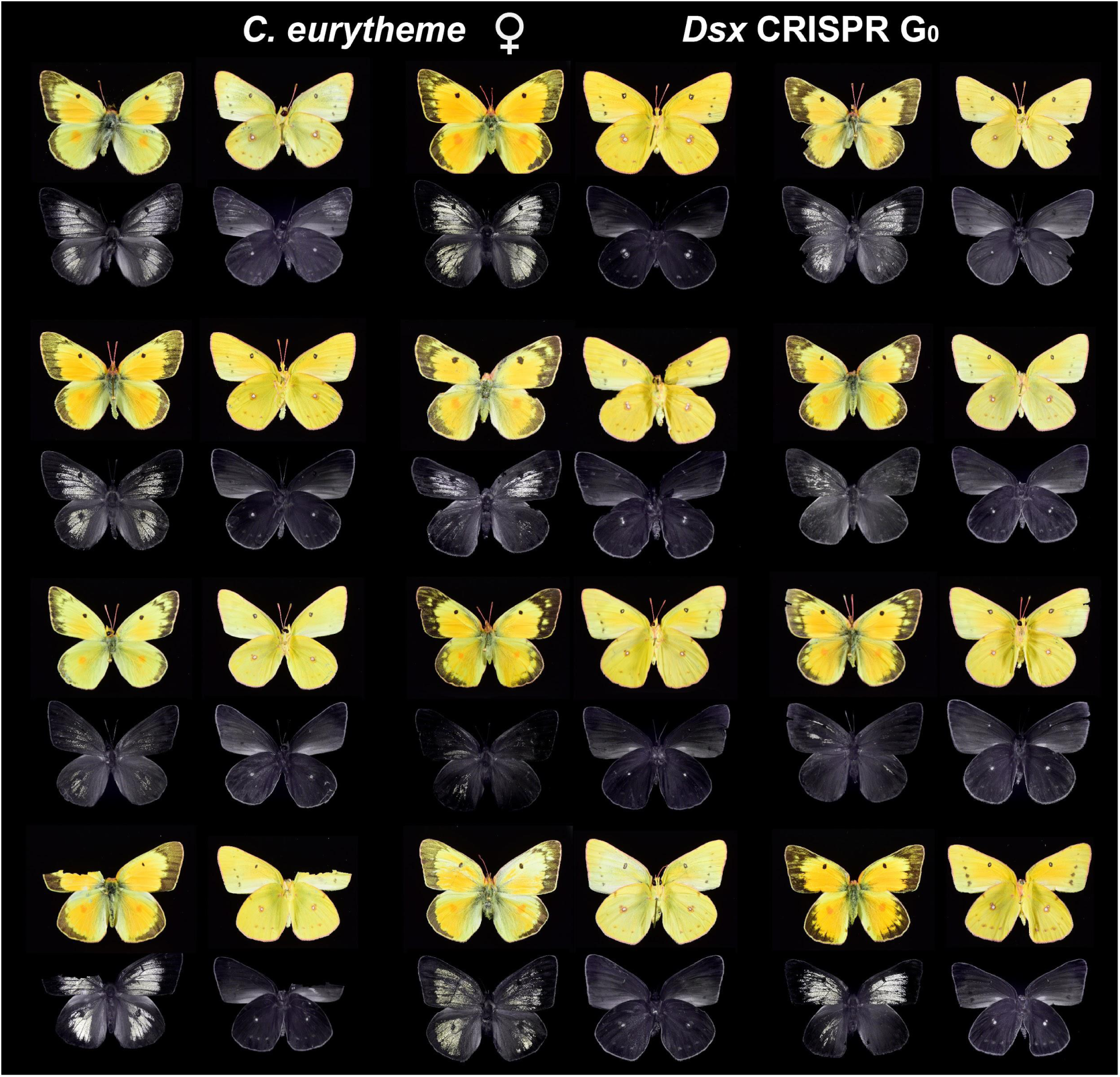
*Dsx* crispant phenotypes in *C. eurytheme* females. Female *Dsx* crispants show a male-like regression of the marginal melanic band, a transformation of marginal melanic scales into male-like scent-related scales (not visible here), and widespread gains of UV-iridescence, all restricted to the dorsal side of each specimen (left panels). There were no visible effects on ventral sides (right panels). Bottom rows : UV-spectrum photography (320-400 nm).

**Figure S4.**
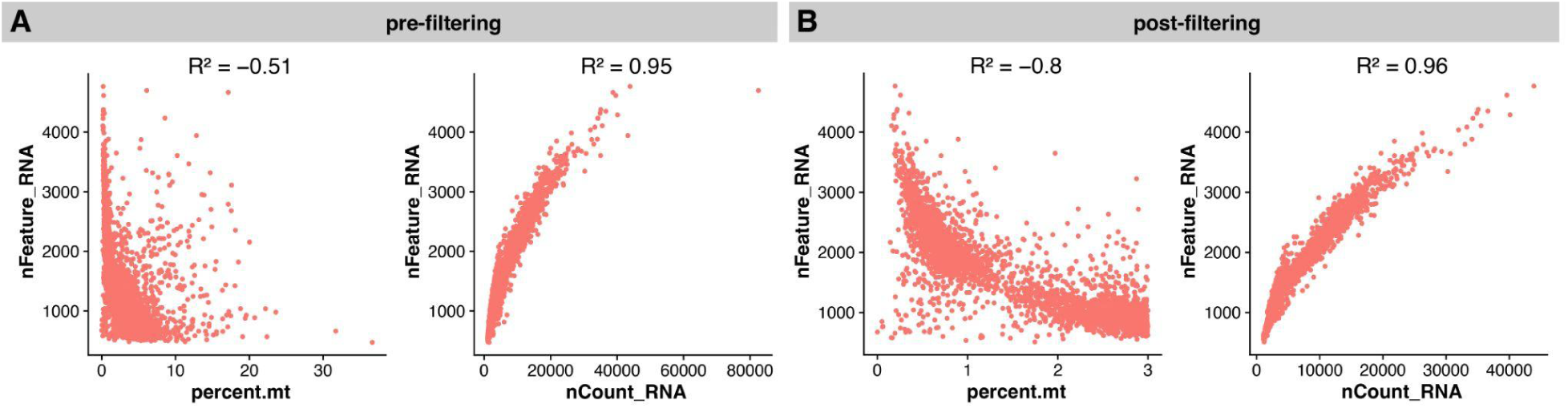
Quality control metrics for 40% *C. eurytheme* wing tissue snRNAseq. These plots compare CellRanger outputs before (A) and after (B) filtering for *nCounts*>3, *nFeatures*>300, *percent.mt*<4%. Left panels : number of genes detected in each cell relative to percentage mitochondrial reads within each cell. Right panels : number of genes detected in each cell relative to the total number of molecules detected within a cell. R^2^ values indicate the coefficient of determination in each comparison.

**Table S1.** Summary of *Dsx* CRISPR KO injections experiments.

| sgRNA | Injected eggs | Injection Time (AEL) | L1 larvae | Pupae | Adults (total) | Adults with Phenotype |
| --- | --- | --- | --- | --- | --- | --- |
| *Ceu_dsx_sgRNA1* | 238 | 1-5h | 43 | 12 | 12 | 2 |
| *Ceu_dsx_sgRNA1* | 96 | 1.5-7h | 15 | 7 | 7 | 1 |
| *Ceu_dsx_sgRNA1* | 250 | 3.5-5h | not recorded | 15 | 18 | 3 |
| *Ceu_dsx_sgRNA2* | 143 | 1-4.5h | 32 | 18 | 18 | 5 |
| *Ceu_dsx_sgRNA2* | 195 | 40min- 4h | 62 | 22 | 22 | 8 |

**Table S2.**
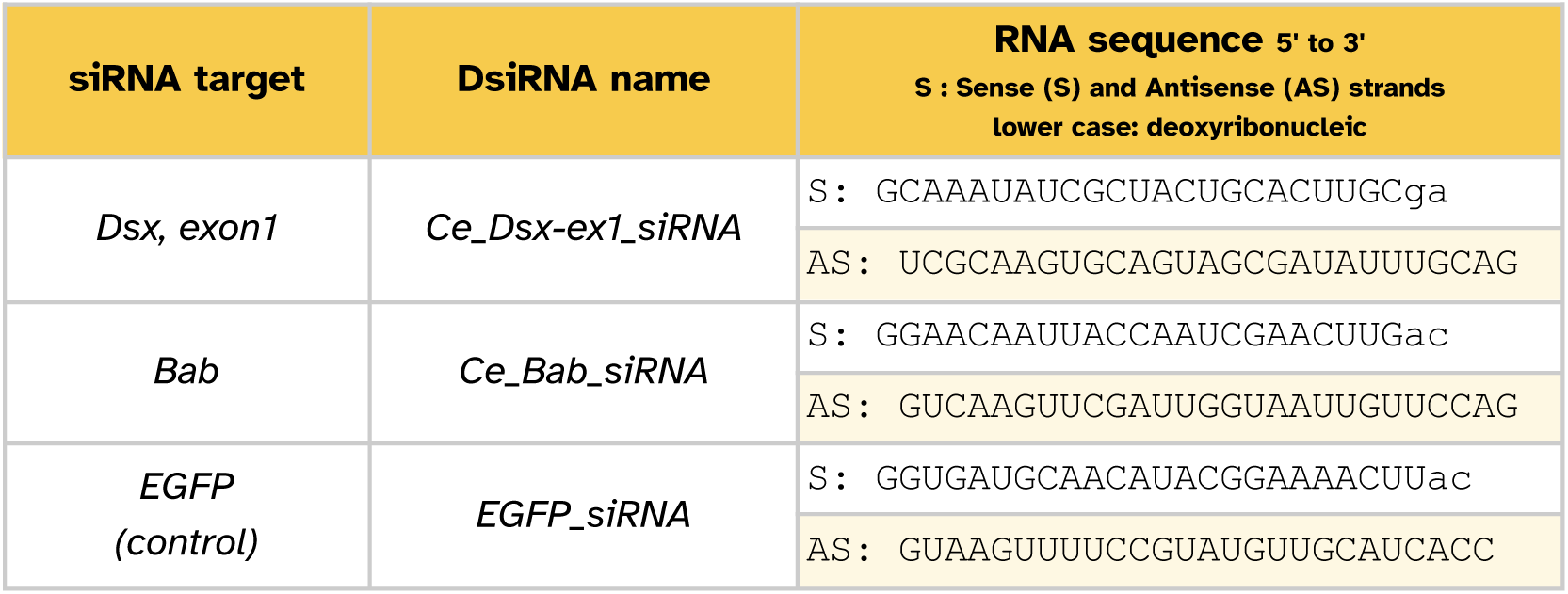
DsiRNA mixes used in electroporation experiments for gene expression knockdowns.

**Data S1. Top differentially expressed genes within each cluster in *C. eurytheme* pupal wing at 40% development.** List of differentially expressed genes (DEGs) marking each cluster using Seurat’s function *FindAllMarkers* (*log2FC* > 0.25, *min.pct* = 0.25, *p* < 0.05, *test.use* = “Wilcox”). Related to **Fig. 6**.

**Data S2. Top differentially expressed genes defining *Scale2+3* from other *Scale* subclusters in *C. eurytheme* pupal wing at 40% development.** List of differentially expressed genes (DEGs) differentiating clusters *Scale2* and *Scale3* from the rest of the clusters using Seurat’s function *FindMarkers* (*log2FC* > 0.25, *min.pct* = 0.25, *p* < 0.05, *test.use* = “Wilcox”). Related to **Fig. 8**.

**Data S3. Top differentially expressed genes defining *Scale2* from *Scale3* in *C. eurytheme* pupal wing at 40% development**

List of differentially expressed genes (DEGs) differentiating *Scale2* from *Scale3* using *Seurat*’s function *FindMarkers* (*log2FC* > 0.25, *min.pct* = 0.25, P < 0.05, *test.use* = “Wilcox”). Related to **Fig. 8**.

**Data S4. Genes with MACS3-called Bab-binding peaks from ChIP-sequencing data from *C. eurytheme* pupal wings at 40% and 60% development**

List of genes with Bab-binding peaks mapped using *MACS3* (*q* value < 0.01) from 2 timepoints, 40% and 60% pupal development with 2 replicates each. Gene list was curated after retaining top N peaks from true replicates below the Irreproducible Discovery Rate (IDR) threshold of 0.05. Related to **Fig. 9**.

**Data S5. Top differentially expressed genes defining *Scale2+3* from other scale subclusters, with Bab-binding peaks in *C. eurytheme* pupal wing at 40% development**

List of genes that are differentially expressed between clusters *Scale2+3* from the rest of the clusters using Seurat’s function *FindMarkers* (*log2FC* > 1.25, *min.pct* = 0.01, adjusted *p* < 0.05, *test.use* = “Wilcox”) and contain Bab-binding peaks at both 40% and 60% development (False Discovery Rate; FDR < 0.05; **Data S4**). Related to **Fig. 9**.

**Data S6. Top differentially expressed genes defining *Scale3* from other scale subclusters with several Bab-binding peaks.** List of 87 genes that are differentially expressed between clusters *Scale3* from the rest of the clusters using Seurat’s function *FindMarkers* (*log2FC* > 1.8, *min.pct* = 0.01, adjusted *p* < 0.01, *test.use* = “Wilcox”) and that contain 3 or more Bab-binding peaks (False Discovery Rate; FDR < 0.05; **Data S4**). Related to **Fig. 10B**.

**Data S7. List of all Bab ChIP sites inside of selective sweeps.** List of 29 genomic positions that contain a *C. eurytheme* selective sweep and a Bab ChIP binding site. Related to **Figs. 10B**, **10D**.

**Data S8. List of genes in proximity to Bab ChIP sites and selective sweeps**. Related to **Fig. 10D**.

